# Neuropilin-1 controls vascular permeability through juxtacrine regulation of endothelial adherens junctions

**DOI:** 10.1101/2024.01.23.576785

**Authors:** Sagnik Pal, Yangyang Su, Emmanuel Nwadozi, Lena Claesson-Welsh, Mark Richards

## Abstract

Neuropilin-1 (NRP1) regulates endothelial cell (EC) biology through modulation of vascular endothelial growth factor receptor 2 (VEGFR2) signalling by presenting VEGFA to VEGFR2. How NRP1 impacts VEGFA-mediated vascular hyperpermeability has however remained unresolved, described as exerting either a positive or a passive function. Using EC-specific *Nrp1* knock-out mice, we discover that EC-expressed NRP1 exerts an organotypic role. In the ear skin, VEGFA/VEGFR2-mediated vascular leakage was increased following loss of EC NRP1, implicating NRP1 in negative regulation of VEGFR2 signalling. In contrast, in the back skin and trachea, loss of EC NRP1 decreased vascular leakage. In accordance, phosphorylation of vascular endothelial (VE)-cadherin was increased in the ear skin but suppressed in the back skin of *Nrp1* iECKO mice. NRP1 expressed on perivascular cells has been shown to impact VEGF-mediated VEGFR2 signalling. Importantly, expression of NRP1 on perivascular cells was more abundant in the ear skin than in the back skin. Global loss of NRP1 resulted in suppressed VEGFA-induced vascular leakage in the ear skin, implicating perivascular NRP1 as a juxtacrine co-receptor of VEGFA in this compartment. Altogether, we demonstrate that perivascular NRP1 is an active participant in EC VEGFA/VEGFR2 signalling and acts as an organotypic modifier of EC biology.

## Introduction

Neuropilin-1 (NRP1) is a multifunctional transmembrane protein that is expressed abundantly on the surface of a range of cell types, where it binds class 3 semaphorins (SEMA3), heparan sulfate and vascular endothelial growth factors (VEGFs) (Goshima et al., 1999, Mamluk et al., 2002, Gu et al., 2002, Soker et al., 1998). VEGFA is the main inducer of VEGF receptor 2 (VEGFR2) signalling in endothelial cells (ECs) and its binding to NRP1 promotes heterocomplex formation between NRP1 and VEGFR2 (Soker et al., 2002). Thus, NRP1, which is devoid of intrinsic catalytic activity, has been designated as a co-receptor of VEGFR2 and is known to modulate VEGFR2 signalling. The cytoplasmic domain of NRP1 contains a C-terminal SEA motif, a PDZ binding domain that mediates binding to synectin (GIPC1) and in-turn regulates the endocytic trafficking of both NRP1 and VEGFR2 (Cai and Reed, 1999, Naccache et al., 2006, Horowitz and Seerapu, 2012, Lanahan et al., 2010). Accordingly, removal of the NRP1 cytoplasmic tail delays VEGFR2 endocytosis following VEGFA binding, leading to enhanced surface retention and reduced phosphorylation of tyrosine (Y)1175 in VEGFR2 (Lanahan et al., 2013). NRP1 thus acts as an important modulator of VEGFR2 activation and its downstream signalling pathways. However, reports have also suggested that VEGFA transduces biological responses in ECs via NRP1, in a VEGFR2-independent manner (Roth et al., 2016).

Global NRP1 knockout in mice results in lethality at E10-E12.5 due to abnormal vessel sprouting in major organs and impaired yolk sac vascularization (Kawasaki et al., 1999). Moreover, NRP1 overexpression leads to excessive vessel growth and promotes leaky and haemorrhagic vessels (Kitsukawa et al., 1995). In contrast, endothelial-specific knockout of NRP1 in mouse embryos only shows mild embryonic brain defects, and in the adult vasculature, no gross abnormalities are evident (Lanahan et al., 2013). Several reports link increased NRP1 expression in tumours to a poor prognosis for survival (Miao et al., 2000, Kawakami et al., 2002, Hong et al., 2007). Furthermore, in diseases such as age related macular degeneration, NRP1 has been linked with increased neovascularization and vessel hyperpermeability, likely due to enhanced signalling downstream of VEGFA (Raimondi et al., 2014, Fernández-Robredo et al., 2017).

VEGFA-induced vascular permeability is mediated through the Y949 and Y1173 phosphorylation sites of VEGFR2. The phosphorylated Y949 presents a binding site for TSAd (T-cell specific adaptor), which in turn binds Src family kinases (SFKs) that phosphorylate and promote internalization of junctional proteins such as VE-cadherin (Matsumoto et al., 2005, Sun et al., 2012, Li et al., 2016, Jin et al., 2022). Concurrently, pY1173 phosphorylation of PLCγ, leading to Ca2+/Protein kinase C (PKC) activation of endothelial nitric oxide synthase (eNOS), contributing to Src activation and phosphorylation of VE-cadherin (Sjoberg et al., 2023).

The role of NRP1 in VEGFA-mediated permeability has been controversial. Studies employing *in vivo* and *in vitro* models have found NRP1 to be a positive regulator of VEGFA-mediated vascular permeability, or to lack effect, dependent on tissue or experimental setup (Fantin et al., 2017, Acevedo et al., 2008, Wang et al., 2015, Becker et al., 2005, Pan et al., 2007, Cerani et al., 2013). Additionally, treatment of ECs with CendR peptides, which bind and induce internalisation of NRP1, increases permeability in a NRP1-dependent, but VEGFR2-independent manner (Roth et al., 2016). Therefore, the exact role of NRP1 in VEGFA-VEGFR2 mediated permeability has remained unclear. NRP1 is a promising target in a number of pathologies where vascular dysfunction is exacerbative. Understanding the relationship between NRP1 and VEGFA-VEGFR2 signalling is thus of potential therapeutic relevance.

Here, we aimed to resolve how NRP1 exerts its effect on VEGFA-mediated vascular permeability. Using mice with global or EC-specific loss of NRP1 expression, we identify a tissue-specific role for endothelial NRP1 in the modulation of VEGFA/VEGFR2 signalling regulating vascular leakage. We find that endothelial-specific loss of NRP1 (*Nrp1* iECKO) can both increase and decrease VEGFA-mediated vascular leakage, dependent on the vascular bed. NRP1 is widely expressed and is an important constituent of perivascular cells in some, but not all, tissues and vessel subtypes. The consequence of global loss of NRP1 expression demonstrates that perivascular NRP1 can modify VEGFA/VEGFR2-induced vascular leakage and steer the tissue-specific role of endothelial NRP1 in barrier integrity. Collectively, these data reveal that the relative expression levels of NRP1 between perivascular cells and ECs acts as a tissue-specific modifier of endothelial VEGFA/VEGFR2 signalling upstream of vascular leakage.

## Results

### NRP1 regulates VEGFA-mediated permeability in an organotypic manner

NRP1 has been studied extensively as a co-receptor of the VEGFA/VEGFR2 pathway, and its ability to modulate VEGFR2 signalling in developmental and pathological angiogenesis has been clearly established. However, results concerning the role of NRP1 in VEGFA-mediated vascular permeability have remained contradictory (Fantin et al., 2017, Acevedo et al., 2008, Wang et al., 2015, Becker et al., 2005, Pan et al., 2007, Cerani et al., 2013). Here, *Nrp1*^fl/fl^; *Cdh5^CreERT2^* (*Nrp1* iECKO) and *Nrp1*^fl/fl^, Cre-negative littermate (Control) mice (Figure 1A) were used to study the EC-specific role of NRP1 in VEGFA-induced vascular leakage (Supplementary Figure 1). A 75% reduction in NRP1 protein levels was achieved in lung tissue, chosen for analysis due to the high EC content (Supplementary Figure 1A). In agreement, immunofluorescent staining and RNA *in-situ* hybridisation (ISH) analyses showed a significant reduction in NRP1 expression in ECs of the ear and back skin (Figure 1B-C, Supplementary Figure 1B-E). Using a highly sensitive assay combining intravital microscopy and atraumatic intradermal microinjection in the ear dermis of *Nrp1* iECKO and *Nrp1^fl/fl^* control mice, VEGFA-induced vascular leakage was significantly increased in venules following the loss of NRP1 in endothelial cells (Figure 1D-E, Video 1-2) (Honkura et al., 2018). Additionally, we measured the rate of dextran extravasation at sites of leakage, which demonstrated that *Nrp1* iECKO mice exhibit more profuse leakage at each site (Figure 1F), indicative of increased disruption of EC-EC junctions. These data were supported by a modified Miles’ assay, with intradermal administration of VEGFA in the ear skin, leading to an increased extravasation of Evans Blue dye in *Nrp1* iECKO mice versus their littermate *Nrp1^fl/fl^* controls (Figure 1G). These findings suggest that NRP1 exerts a stabilizing role by suppressing VEGFA-induced endothelial junction disruption in the ear dermis. This finding is in contrast to reports showing NRP1 to be a positive regulator of VEGFA-mediated vascular leakage (Fantin et al., 2017, Acevedo et al., 2008, Becker et al., 2005).

**Figure 1:**
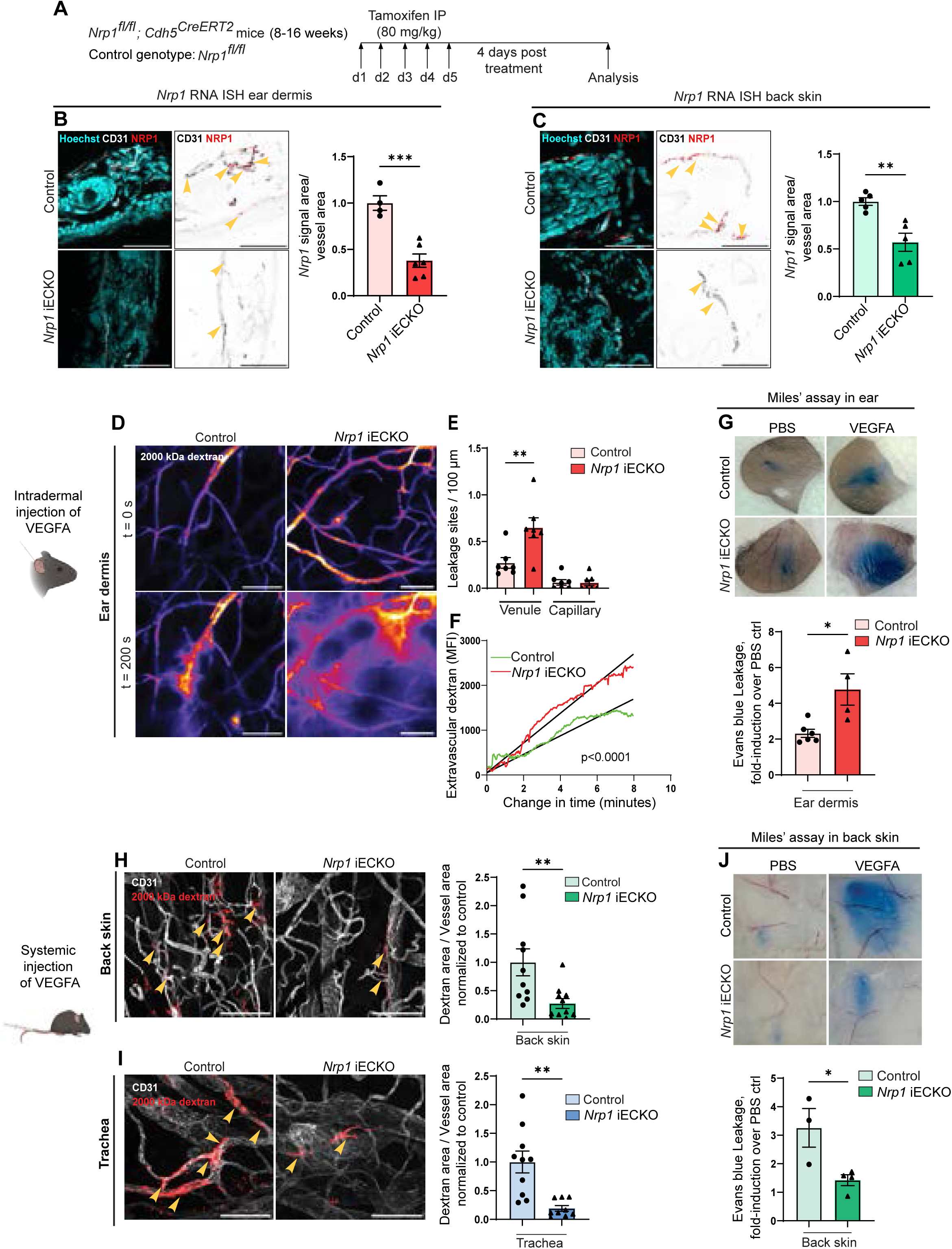
NRP1 regulates VEGFA-mediated permeability in an organotypic manner. **(A)** Schematic illustration of experimental design to induce recombination in *Nrp1^fl/fl^; Cdh5^CreERT2^* (*Nrp1* iECKO) mice. *Nrp1^fl/fl^*were used as control. **(B-C)** RNA *in-situ* hybridisation (ISH) analysis of *Nrp1* expression in the ear dermis **(B)** and back skin **(C)** of *Nrp1* iECKO mice. Images highlight *Nrp1* mRNA particles (arrows) specific to CD31-positive vessel area and graphs show quantification of *Nrp1* mRNA expression (*Nrp1* mRNA particle area / vessel area). Scale bar: 50 μm. **(D)** Representative images showing leakage of 2000 kDa FITC-dextran (Pseudo-colour) in response to intradermal VEGFA injection in ear dermis of control and *Nrp1* iECKO mice. **(E)** Leakage sites per vessel length in response to intradermal VEGFA stimulation in the ear skin of control and *Nrp1* iECKO mice. n=6 mice, two or more acquisitions / mouse. **(F)** Quantification of 2000 kDa dextran extravasation over time in the ear skin of control and *Nrp1* iECKO mice following intradermal VEGFA stimulation. Black lines represent lines of best fit for the slope between leakage initiation and leakage termination. N ≥ 3 mice, two or more acquisitions / mouse, three or more sites / acquisition. **(G)** Evans blue leakage following intradermal administration of VEGFA in the ear skin of control and *Nrp1* iECKO mice. Top, representative images. Bottom, quantification of Evans blue extravasation shown as VEGFA-induced leakage fold over PBS control (n ≥ 4 mice). **(H-I)** Leakage of fixable 2000 kDa FITC dextran in back skin **(H)** and trachea **(I)** after systemic administration of VEGFA in control and *Nrp1* iECKO mice Left, representative images. Right, quantification of tracer leakage area / vessel area (n ≥ 8 mice, 2 or more fields of view / mouse). **(J)** Evans blue leakage with intradermal administration of VEGFA in the back skin of control and *Nrp1* iECKO mice. Top, representative images. Bottom, quantification of Evans blue extravasation shown as VEGFA-induced leakage fold over PBS control (n ≥ 4 mice). Error bars; mean ± SEM. Statistical significance: Two-tailed unpaired Student’s t-test and linear regression with ANCOVA. Scale bar: 100 μm unless stated.

One possible explanation for these differences compared to previous reports is that the role of NRP1 may be organ dependent, in line with the growing insights into the distinct properties of ECs in different vessel types and vascular beds (Richards et al., 2022, Richards et al., 2021, Augustin and Koh, 2017). To investigate a potential organotypic role for NRP1 in EC biology, we studied the consequence of *Nrp1* knockout on the barrier properties of ECs in different vascular beds.

*Nrp1* iECKO and their littermate controls were assessed, first for their basal permeability properties, revealing that loss of EC NRP1 did not significantly alter basal permeability of a 10 kDa dextran tracer (Supplementary Figure 2A). To asses VEGFA-induced vascular leakage, selected organs were initially assessed for their susceptibility to VEGFA, chosen based on prior experience of different EC barrier properties (Richards et al., 2022, Richards et al., 2021, Aird, 2007, Augustin and Koh, 2017). Mice were challenged with systemic administration of fluorescent dextrans with or without VEGFA for 30 minutes before collecting organs and assessing acute vascular leakage microscopically or following solvent-based extraction of dextran and fluorescence spectroscopy. VEGFA-induced vascular leakage was observed in back skin, trachea, kidney, skeletal muscle and heart (Supplementary Figure 2B-C). Subsequently, VEGFA-induced vascular leakage was assessed in *Nrp1* iECKO tissues, demonstating that loss of NRP1 decreased VEGFA-mediated vascular leakage in the trachea and back skin (Figure 1H-I), but had no effect on kidney, skeletal muscle and heart irrespective of tracer size (70 kDa and 2000 kDa; Supplementary Figure 2D-E).

Using a Miles’ assay, endothelial cell NRP1’s positive regulation of VEGFA-VEGFR2 signalling in the back skin was confirmed, with a reduction in VEGFA-induced vascular leakage in *Nrp1* iECKO mice (Figure 1J), in keeping with previous publications (Fantin et al., 2017). These data collectively suggest that endothelial NRP1 has an organ-specific role in controlling VEGFA-mediated vascular permeability. Importantly, endothelial NRP1 is a positive regulator of VEGFA mediated permeability in the trachea and back skin but a negative regulator in the ear skin.

### Global inactivation of NRP1 reduces VEGFA-mediated vascular permeability

Our results here, where NRP1 may play positive (trachea and back skin), negative (ear skin) and passive (kidney, skeletal muscle and heart) roles in VEGFA-induced vascular leakage, were collected using EC-specific knockout mice. In previous studies, mouse models of both EC-specific deletion and a globally expressed NRP1 C-terminal deletion mutant were employed (Fantin et al., 2017, Roth et al., 2016). We thus set out to investigate whether global inactivation of NRP1 might differently modify the VEGFA-induced leakage response.

*Nrp1^fl/fl^* mice were crossed with *Actb^Cre^* mice to generate global *Nrp1* knock-out mice (*Nrp1* iKO) for which *Nrp1^fl/fl^* mice were used as controls. Efficient reduction of NRP1 protein in the *Nrp1* iKO was confirmed in lung lysates (Figure 2A-B). Complete loss of NRP1 could also be seen in the ear and back skin of these mice using immunofluorescent staining (Supplementary Figure 1A and B). Intravital imaging of the ear dermis of *Nrp1* iKO mice showed that global loss of NRP1 suppressed VEGFA-induced vascular leakage (Figure 2C-D, Video 3-4), in contrast to the increased leakage seen in *Nrp1* iECKO (see Figure 1D-G). Furthermore, global *Nrp1* knock-out resulted in a ∼75% reduction in VEGFA-mediated vascular leakage also in the back skin and tracheal vasculatures, similar to that seen in *Nrp1* iECKO mice (Figure 2E-F). Thus, the global loss of NRP1 results in an alignment of leakage phenotype across the studied tissues.

**Figure 2:**
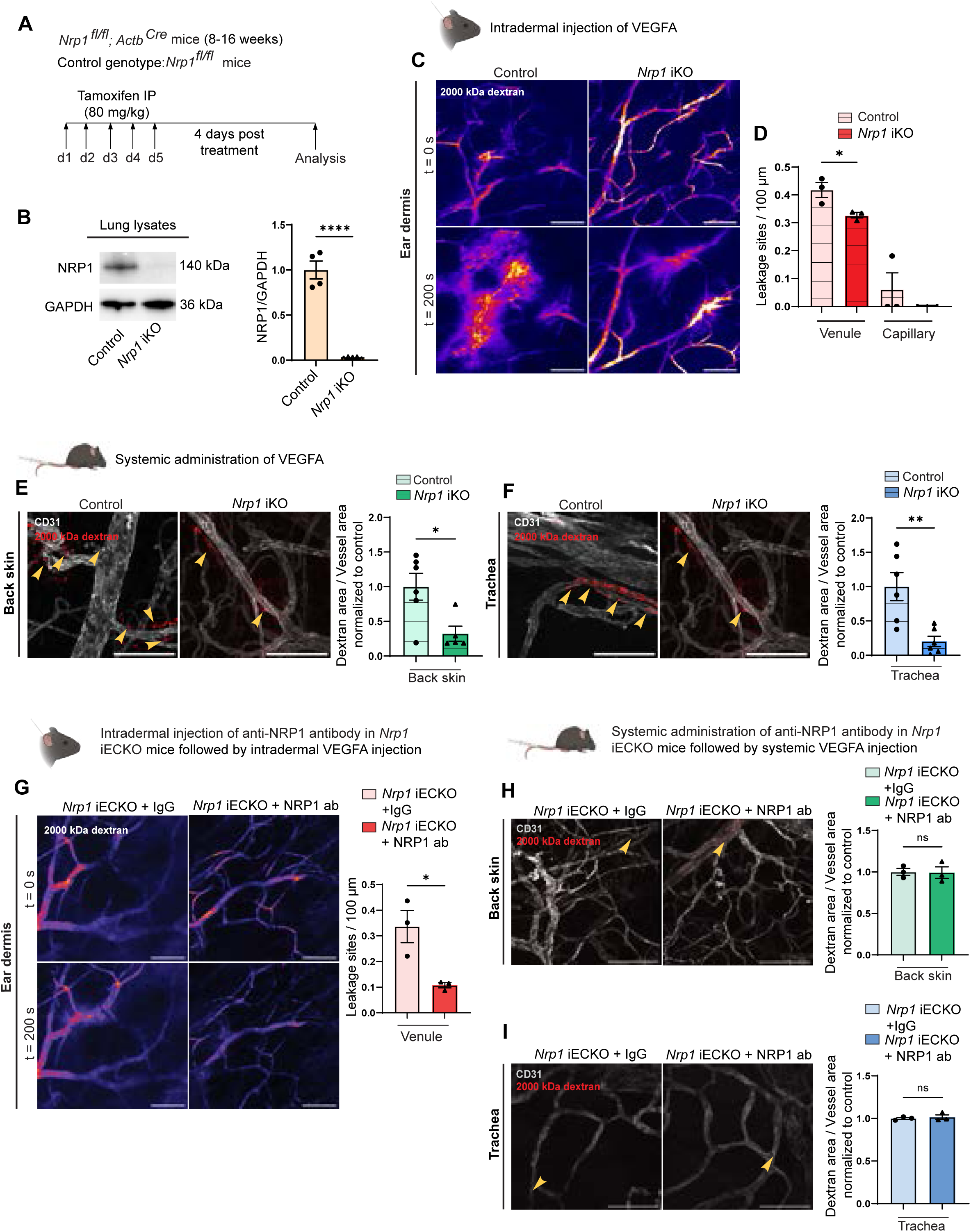
Global loss of NRP1 reduces VEGFA-mediated vascular permeability. **(A)** Schematic illustration of experimental design to induce recombination in *Nrp1^fl/fl^; Actb^Cre^* (*Nrp1* iKO) mice. *Nrp1^fl/fl^* were used as control. **(B)** Western blot and quantification of NRP1 protein levels in lung lysates from tamoxifen-treated control and *Nrp1* iKO mice (n ≥ 3 mice). **(C)** Representative images showing leakage of 2000 kDa FITC dextran (Pseudo-colour) in response to intradermal VEGFA injection in the ear of control and *Nrp1* iKO mice. **(D)** Leakage sites per vessel length in response to intradermal VEGFA stimulation in the ear skin of control and *Nrp1* iKO mice. N = 3 mice, two or more acquisitions / mouse. **(E-F)** Leakage of fixable 2000 kDa FITC dextran in back skin **(E)** and trachea **(F)** after systemic administration of VEGFA in control and *Nrp1* iKO mice. Left, representative images. Right, quantification of tracer leakage area / vessel area (n ≥ 5 mice, 3 or more fields of view / mouse). **(G)** VEGFA-induced leakage in the ear skin of *Nrp1* iECKO mice treated intradermally with isotype control or NRP1-VEGFA blocking antibody. Left, representative images. Right, quantification of leakage sites per 100 µm of vessel length (n = 3 mice, ≥ 2 acquisitions / mouse). **(H-I)** VEGFA-induced leakage of 2000 kDa dextran in back skin **(H)** and trachea **(I)** of *Nrp1* iECKO mice treated systemically with isotype control or NRP1-VEGFA blocking antibody. Left, representative images. Right, quantification of tracer area / vessel area (n = 3 mice). Error bars; mean ± SEM. Statistical significance: Two-tailed unpaired Student’s t-test. Scale bar: 100 μm.

These conclusions were further explored through the use of an antibody that blocks the binding of VEGFA to NRP1 (Termini et al., 2021). Administration of this blocking antibody in *Nrp1* iECKO mice, where only endothelial cell NRP1 is removed, resulted in suppressed VEGFA-induced vascular leakage in the ear dermis, while vascular leakage was unchanged in the back skin and trachea (Figure 2G-I, Video 5-6).

These data illustrate that NRP1 modulates VEGFA signaling in an organotypic and cell-specific manner. Notably, we demonstrate that VEGFA-induced leakage in the back skin and trachea is indistinguishable between global and endothelial-specific NRP1 loss, while there is a contrasting phenotype in the ear skin.

### NRP1 is heterogeneously expressed in perivascular cells

NRP1 is known to form a heterocomplex with VEGFR2 upon VEGFA binding. Interestingly, VEGFR2/NRP1/VEGFA can assemble as a juxtacrine *trans* complex, where NRP1 and VEGFR2 are expressed on the surface of adjacent cells, as well as a *cis* complex, where NRP1 and VEGFR2 are expressed on the same cell (Koch et al., 2014). Importantly, even though the kinetics of *trans* VEGFR2/NRP1 complex formation is slow, these complexes are stable and produce a distinct signalling output compared to the *cis* configuration (Koch et al., 2014). Thus, NRP1 presented in *cis* or *trans* produces differential VEGFR2 signalling output upon VEGFA binding. Given the above findings we reasoned that peri-endothelial distribution of NRP1 could be an important modifier of EC VEGFR2 signalling and possibly explain organotypic differences observed in *Nrp1* iECKO mice.

To investigate the pattern of NRP1 expression we employed a *Pdgfrβ* promoter-driven GFP (*Pdgfrβ-EGFP*) reporter mouse to visualize PDGFRβ-positive perivascular cells. To visualize ECs, and the localization of NRP1 in the ear skin and back skin from these mice, tissues were immunostained for CD31 and NRP1. NRP1 expression could be seen in both endothelial and perivascular cells, which was lost in both cell types after global *Nrp1* deletion in *Nrp1* iKO mice, while its expression remained expressed in perivascular cells in *Nrp1* iECKO mice (Figure 3A and Supplementary Figure 1B-E). Both arteriolar (Figure 3B) and capillary (Figure 3C) PDGFRβ-positive perivascular cells expressed NRP1. The relative NRP1 expression in perivascular cells compared to EC was consistently higher in the ear skin compared to back skin. Strikingly, the low perivascular NRP1 expression in the back skin resulted in a 4-fold difference in the perivascular/EC NRP1 ratio when comparing venules of the back skin with the ear skin (Figure 3D).

**Figure 3:**
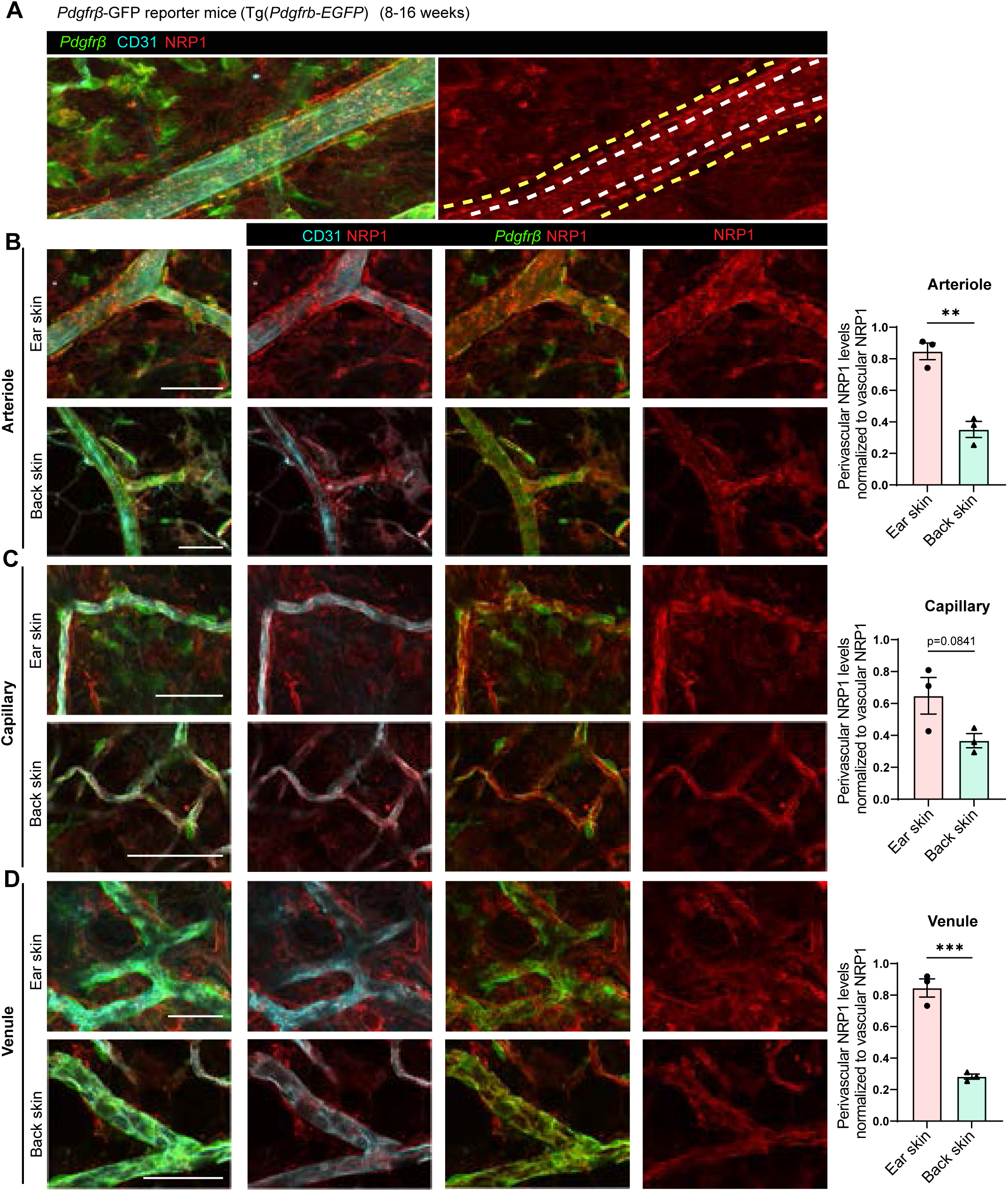
Perivascular expression of NRP1 is heterogeneous. **(A)** Representation of NRP1 expression and method of NRP1 quantification in endothelial and perivascular cells of *Pdgfrβ-*GFP mice. White dashed lines in right image outlines vascular area, space between white and yellow dashed lines represents perivascular area. Note perivascular, NRP1 expressing cells in between yellow and white lines. **(B-D)** Images, left, and quantification, right, showing vascular and perivascular NRP1 expression in arterioles **(B)**, capillaries **(C)** and venules **(D)** of ear skin and back skin and its relative expression. Error bars; mean ± SEM. Statistical significance: Two-tailed unpaired Student’s t-test. Scale bar: 50 μm.

These data thus show that NRP1 is expressed in perivascular cells of the ear skin and back skin. We find however that the ratio of NRP1 expression between ECs and perivascular cells differs between ear skin and back skin, and between different vessel types. In ear skin the perivascular/EC NRP1 ratio is higher compared to the back skin, which may support a higher ratio of *trans* NRP1/VEGFR2 relative to cis complexes in the ear skin.

### NRP1 distribution modifies VEGFA-mediated signalling and vascular leakage

NRP1 modifies VEGFR2 signalling by controlling its internalisation and intracellular trafficking (Bayliss et al., 2020, Ballmer-Hofer et al., 2011). The presence of NRP1 in *trans* however, modifies this dynamic by retaining VEGFR2 on the cell surface for longer time periods, altering its signalling output (Koch et al., 2014).

We thus wished to investigate whether perivascular NRP1 expression might impact VEGFR2 signalling upstream of vascular leakage, and explain the above described organotypic effects of EC NRP1 loss. For this purpose, *Nrp1* iECKO mice were crossed with homozygous *Vegfr2^Y949F/Y949F^* mice, to produce *Vegfr2^Y949F/Y949F^*;*Nrp1* iECKO mice that are deficient in both EC NRP1 expression and VEGFR2 signalling upstream of vascular leakage. In these mice, loss of NRP1 protein was efficiently established in lung tissue (Figure 4A-B). Analysis of VEGFA-mediated vascular leakage in the ear dermis showed that the increased leakage, induced by the loss of EC NRP1, was abrogated by the concurrent loss of the VEGFR2 phosphosite Y949, known to be required for activation of the cytoplasmic tyrosine kinase Src in response to VEGFA (Figure 4C-D, Video 7-8). These analyses suggest that, in the ear dermis, EC NRP1 negatively regulates VEGFA-induced vascular leakage by modulation of VEGFR2 activation and phosphorylation.

**Figure 4:**
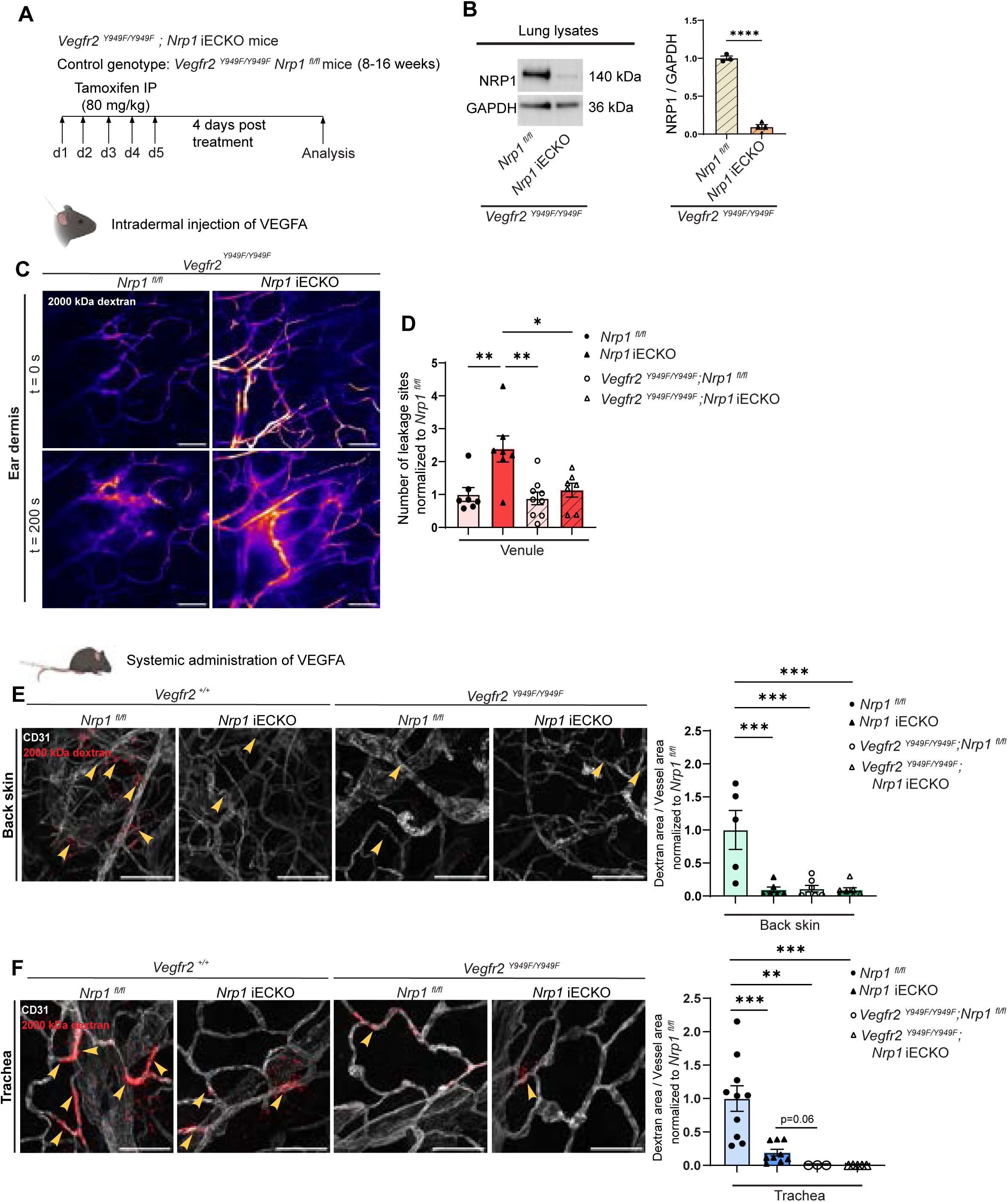
NRP1 distribution modifies VEGFA-induced vascular permeability. **(A)** Schematic illustration of experimental design to induce recombination in *Vegfr2 ^Y949F/Y949F^*; *Nrp1* iECKO mice. *Vegfr2 ^Y949F/Y949F^*; *Nrp1^fl/fl^* mice were used as control. **(B)** Western blot and quantification of NRP1 using lung lysates from tamoxifen-treated control and *Vegfr2 ^Y949F/Y949F^*; *Nrp1* iECKO mice (n = 3 mice). **(C)** Representative images showing leakage in response to intradermal VEGFA injection in the ear of control *Vegfr2 ^Y949F/Y949F^*; *Nrp1^fl/fl^*mice and *Vegfr2 ^Y949F/Y949F^*; *Nrp1* iECKO mice. **(D)** Leakage sites per vessel length in response to intradermal VEGFA stimulation in the ear skin of control *Vegfr2^Y949F/Y949F^*; *Nrp1^fl/fl^* and *Vegfr2^Y949F/Y949F^*; *Nrp1* iECKO mice. Note, data in *Nrp1^fl/fl^* and *Nrp1* iECKO mice from Figure 1E are also shown for comparative purposes. Data are normalized to appropriate control (n ≥ 6 mice, two or more acquisitions / mouse). **(E-F)** Leakage of 2000 kDa dextran in back skin **(E)** and trachea **(F)** of *Nrp1^fl/fl^*, *Nrp1* iECKO, *Vegfr2 ^Y949F/Y949F^* and *Vegfr2^Y949F/Y949F^*; *Nrp1* iECKO mice after systemic administration of VEGFA. Left, representative images. Right, quantification of tracer leakage area/vessel area (n ≥ 3 mice, 2 or more fields of view / mouse). Error bars; mean ± SEM. Statistical significance: Two tailed unpaired Student’s t-test and Two-way ANOVA. Scale bar: 100 μm.

We next sought to determine the relationship between NRP1 and VEGFR2 Y949 signalling in the trachea and back skin vasculatures. Leakage in *Vegfr2^Y949F/Y949F^*;*Nrp1* iECKO mice was induced by systemic VEGFA and fluorescent 2000 kDa dextran administration in both *Nrp1* iECKO and *Vegfr2^Y949F/Y949F^*;*Nrp1* iECKO mice, and their appropriate controls. As we observed previously, loss of EC NRP1 reduced vascular leakage in the back skin and trachea by 75%, an effect that was enhanced further by the VEGFR2 Y949F mutation in tracheal vasculature (Figure 4E-F). Importantly, leakage in trachea and back skin was similarly inhibited in the *Vegfr2^Y949F/Y949F^*model as in the combined *Vegfr2^Y949F/Y949F^*;*Nrp1* iECKO model, revealing that NRP1’s positive regulation of vascular leakage in these tissues is mediated through VEGFR2 Y949. Altogether, these data show that NRP1 presented in *trans* by perivascular cells in the ear dermis can signal through VEGFR2 to induce vascular leakage, and that VEGFR2 Y949 is vital for mediating NRP1’s effects on endothelial junctions.

### Perivascular NRP1 modifies VEGFA-induced signalling

VEGFA/VEGFR2 signalling mediates Src activation and phosphorylation of VE-Cadherin on Y685, required for vascular permeability (Orsenigo et al., 2012, Wessel et al., 2014). We have also recently shown that PLCγ is an important mediator of VEGFA-induced vascular leakage by allowing the production of nitric oxide through eNOS, which in turn modified Src by tyrosine nitration, required for full Src activity (Sjoberg et al., 2023). To clarify the impact of perivascular NRP1 on VEGFA-induced signalling we studied VE-Cadherin and PLCγ phosphorylation in *Nrp1* iECKO and *Nrp1* iKO mice. Perivascular NRP1 expression present in the ear dermis of *Nrp1* iECKO mice was lost in *Nrp1* iKO mice (Supplementary Figure 1B-C). In agreement with the enhanced vascular leakage in the ear dermis of *Nrp1* iECKO mice (Figure 1D-G), VEGFA-induced VE-Cadherin phosphorylation at Y685 was enhanced in *Nrp1* iECKO mice versus their littermate control (Figure 5A). No such induction however was seen in *Nrp1* iKO mice (Figure 5B). In the back skin, VE-Cadherin pY685 levels were induced by VEGFA in the control but not in the *Nrp1* iECKO, nor the iKO models (Figure 5C-D), in accordance with the described permeability phenotype (see Figure 1, panels H-I). Similarly, PLCγ phosphorylation was potentiated in the ear skin of *Nrp1* iECKO but not *Nrp1* iKO mice (Supplementary Figure 3A-B), whereas in the back skin, loss of either endothelial or global NRP1 led to a reduction in PLCγ phosphorylation following intradermal VEGFA stimulation (Supplementary Figure 3C-D).

**Figure 5:**
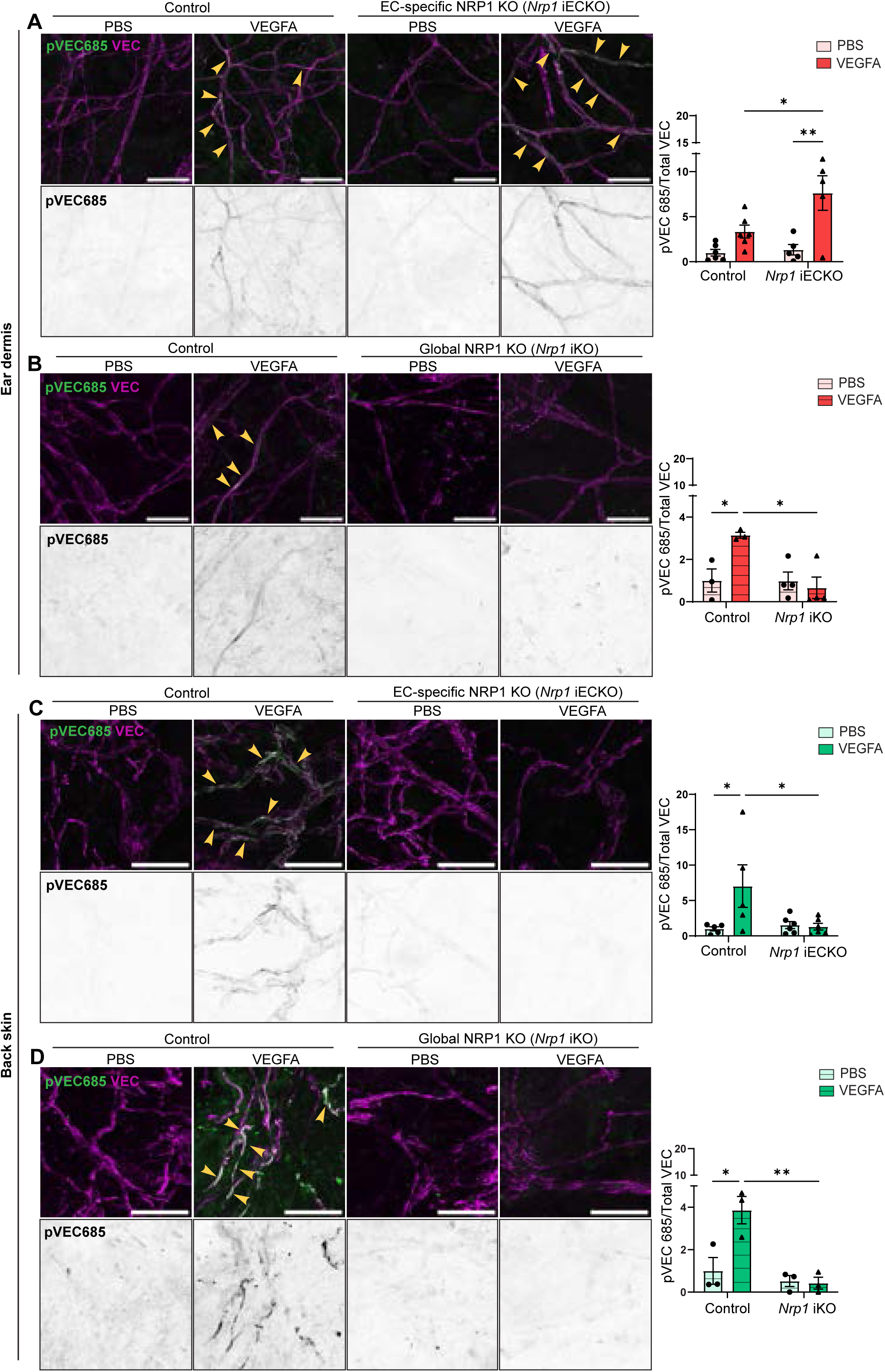
Perivascular NRP1 modifies VEGFA-induced signalling. **(A-B)** Phosphorylation of VE-Cadherin (VEC) Y685 in response to intradermal PBS or VEGFA injections in the ear dermis of control *Nrp1^fl/fl^* and *Nrp1* iECKO **(A)** or *Nrp1* iKO **(B)** mice. Left, representative images. Right, quantification of phosphorylated VEC Y685 (pVEC685) area per total VEC area after VEGFA stimulation, normalized to PBS in the ear dermis of control *Nrp1^fl/f^* and *Nrp1* iECKO or *Nrp1* iKO mice. N ≥ 3 mice, two or more fields of view / mouse. **(C-D)** Phosphorylation of VE-Cadherin Y685 in response to intradermal PBS or VEGFA injections in the back skin of control *Nrp1^fl/f^* and *Nrp1* iECKO **(C)** or *Nrp1* iKO **(D)** mice. Left, representative images. Right, quantification of phosphorylated VEC Y685 (pVEC685) area per total VEC area after VEGFA stimulation, normalized to PBS, in the back skin of control *Nrp1^fl/f^* and *Nrp1* iECKO or *Nrp1* iKO mice. N ≥ 3 mice, two or more fields of view / mouse. Error bars; mean ± SEM. Statistical significance: Two-way ANOVA. Scale bar: 50 μm.

Taken together these data show that NRP1 is expressed by perivascular cells in a tissue-specific manner and that perivascular NRP1 can be an important modulator of VEGFA/VEGFR2 signalling. In the ear dermis, NRP1 is expressed by both endothelial and perivascular cells, forming *cis* and *trans* interactions with VEGFR2, respectively. Whilst the *cis* EC interaction predominates, loss of EC NRP1 promotes *trans* NRP1/VEGFR2 interaction and potentiates VEGFA-induced signalling regulating vascular leakage (Figure 6). Meanwhile, due to a paucity of perivascular NRP1 in the trachea and back skin vasculatures, loss of EC NRP1 leads to a loss of NRP1:VEGFR2 complexes, attenuating VEGFA-regulated vascular leakage.

**Figure 6:**
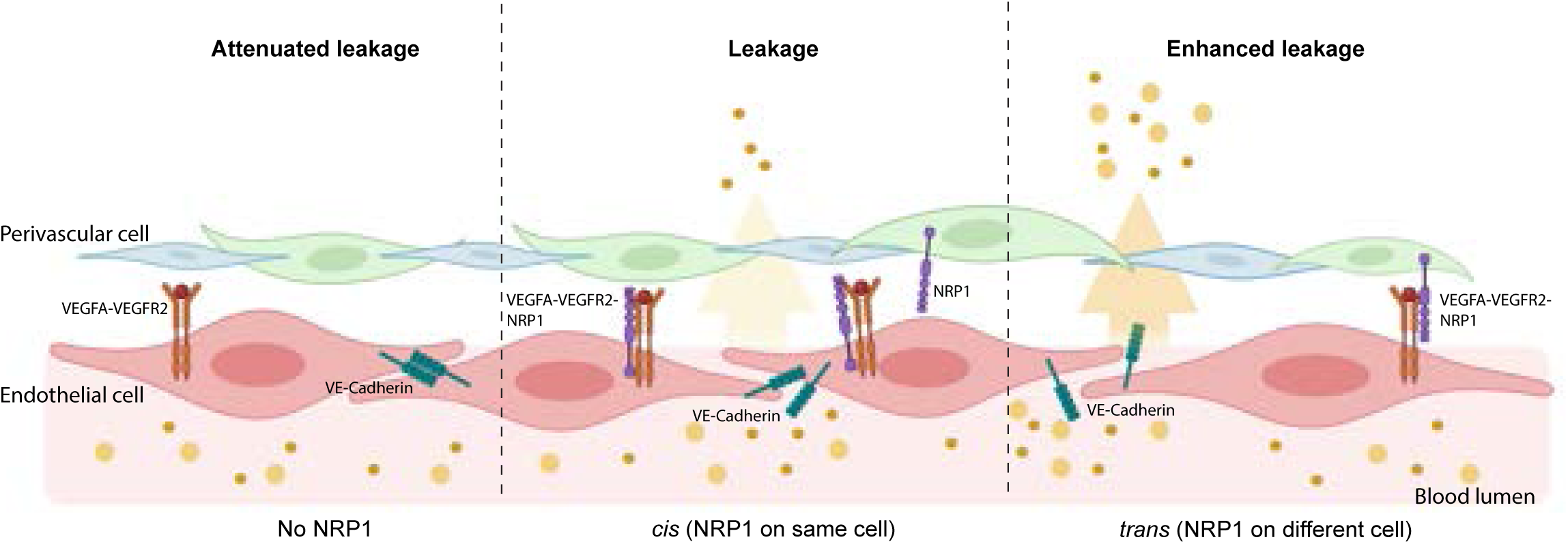
Schematic model of NRP1’s spatial effects on VEGFA-induced vascular permeability. With global loss of NRP1, VEGFA/VEGFR2 signalling regulating vascular leakage is suppressed and thus leakage is reduced (left). In the presence of EC NRP1, a VEGFA/VEGFR2/NRP1 complex is formed in *cis,* which has a rapid and transient effect on VEGFR2 downstream signalling (middle). Loss of EC NRP1 but the presence of perivascular NRP1 leads to formation of a stable VEGFA/VEGFR2/NRP1 complex in *trans* and enhanced VEGFR2 signalling and increased vascular permeability.

## Discussion

Here we find that NRP1 expression by perivascular cells is an important modulator of VEGFA signalling regulating endothelial junction stability. Our data suggest that NRP1 presented in a juxtacrine manner can potentiate VEGFR2 signalling upstream of vascular leakage. NRP1 is expressed by a number of cells adjacent to blood vessels, including PDGFRβ-positive pericytes. However, the abundance of these cells and their level of NRP1 expression differ between vascular beds. Loss of EC-specific NRP1 thus has a differential effect on VEGFA-mediated vascular leakage in different vascular beds, dependent on the ability of ECs to form *tran*s NRP1/VEGFR2 complexes.

Numerous studies have implicated endothelial NRP1 as a positive regulator of permeability (Fantin et al., 2017, Becker et al., 2005, Acevedo et al., 2008). However, multiple studies argue against this, with NRP1 blockade or deletion having no effect on VEGFA-induced vascular leakage in the skin and retina (Pan et al., 2007, Cerani et al., 2013). Here we find that the function of endothelial NRP1 in VEGFA-mediated vascular permeability varies in different vascular beds. In keeping with previous data, we find that loss of NRP1 reduces VEGFA-induced vascular leakage in the back skin and trachea. Meanwhile, endothelial NRP1 appears to have no effect on VEGFA-induced vascular leakage in kidney, skeletal muscle and heart. In the ear skin, however, loss of NRP1 surprisingly leads to an increase in VEGFA-induced vascular leakage. This appears to be through enhanced VEGFR2 Y949 signalling, as mutation of this phosphosite normalises the enhanced leakage seen after loss of NRP1. Previous publications have described NRP1 as being pivotal for VEGFR2-induced Src activity via recruitment of Abl kinase (Fantin et al., 2017). How exactly VEGFA-VEGFR2 signals to Abl, and whether this requires the Y949 phosphosite, however is unknown. Further work is required to understand exactly how NRP1 modifies VEGFR2 activity and how this differs between different vascular beds.

As others have noted and we see here, NRP1 is expressed by perivascular cells, including PDGFRβ positive pericytes (Bartlett et al., 2017, Wnuk et al., 2017). However, we find that perivascular NRP1 expression and its ratio to EC expression differs between different vascular beds, being higher in the ear skin than in back skin. When the *trans* NRP1/VEGFR2 complex predominates, as in the ear skin of *Nrp1* iECKO mice, our data indicate that VEGFA/VEGFR2 signalling impacting junction stability is potentiated. This is in keeping with reports showing that NRP1 presented in *trans* can modify VEGFR2 signalling kinetics, resulting in prolonged PLCγ and ERK2 phosphorylation (Koch et al., 2014). However how such prolonged kinetics might affect the VEGFR2 signalling pathways upstream of vascular leakage has not been previously explored. In addition, juxtacrine NRP1 interactions have been shown to have important functions in cell types other than ECs. For example, NRP1 expressed on microglia is important for the trans-activation of PDGFRα on oligodendrocyte precursor cells, resulting in enhanced proliferation and myelin repair (Sherafat et al., 2021). Furthermore, NRP1 has been posited as a potential immune checkpoint inhibitor (Sarris et al., 2008). NRP1 is highly expressed by regulatory T cells, as well as antigen presenting dendritic cells, and the formation of *tran*s NRP1 homodimers has been suggested to stabilise regulatory T cell function and promote immune tolerance. Finally, numerous studies have described the expression of NRP1 on tumour cells, which can modify EC behaviour and alter their angiogenic potential (Koch et al., 2014, Miao et al., 2000, Hu et al., 2007, Shahrabi-Farahani et al., 2016), with an impact on tumour cell proliferation in human pancreatic and renal cell cancer (Koch et al., 2014, Morin et al., 2020, Morin et al., 2018). This may at least in part be due to the ability of NRP1 to bind VEGFA and thus provide a pool of tumoural VEGFA towards which ECs migrate.

One possible mechanism by which NRP1 presented in *trans* might alter VEGFA/VEGFR2 signalling, when compared to *cis*, is via modification of VEGFR2 trafficking. The adapter molecule synectin is an important intracellular binding partner of NRP1 and facilitates myosin IV-mediated endocytosis (Cai and Reed, 1999, Naccache et al., 2006). A recent study also found that NRP1 interacts with members of the ERM family of intracellular proteins, which act as intermediaries between the plasma membrane and actin cytoskeleton and that have also been associated with endocytosis and intracellular trafficking (Ponuwei, 2016, Gioelli et al., 2022). NRP1 may thus function not just as a co-receptor, but also as a linker to mediate the intracellular trafficking of cell surface receptors. This idea is supported by studies showing that NRP1 co-ordinates PDGFRα internalisation and its intracellular signalling, as well as the trafficking of integrin α5β1 during cell adhesion (McGowan and McCoy, 2021). Furthermore, it has recently been shown that NRP1 binds to VE-Cadherin and regulates its cell surface turnover, a role that has been proposed to at least partly explain its role in EC barrier integrity (Gioelli et al., 2022, Bosseboeuf et al., 2023). In response to VEGFA, the cytoplasmic domain of NRP1, which binds synectin, has been found to be crucial for NRP1’s positive regulation of leakage (Fantin et al., 2017). Furthermore, mutant mice lacking the NRP1 cytoplasmic domain have impaired arteriogenesis due to delayed VEGFR2 endocytosis and resulting reduction in phosphorylation at Y1175 (Lanahan et al., 2013). Interestingly, the C-terminal tail is also crucial for the stability of trans NRP1-VEGFR2 complexes (Koch et al., 2014). A central function of NRP1 is thus to mediate VEGFR2 internalisation and trafficking following VEGFA stimulation. From our data we hypothesise that NRP1 presented in *trans* delays VEGFR2 internalisation and potentiates VEGFR2 signalling upstream of vascular leakage. Indeed, a recent study found that internalisation of VEGFR2 is required to shutdown VEGFR2 Y949 signalling through increasing the stoichiometric ratio of cell surface phosphatases that dephosphorylate Y949 (Corti et al., 2022). Our data is compatible with the hypothesis that VEGFR2 held at the cell surface via juxtracrine NRP1 interaction is capable of enhancing VEGFR2 Y949 signalling, resulting in the disruption of EC junctions. However, due to the rapid turnover of VEGFR2 phosphorylation, this is technically very challenging to conclusively demonstrate. We hypothesize however that NRP1 expression within the perivascular microenvironment could be an important modifying factor of EC signalling and function.

Combined, we demonstrate that different levels and patterning of perivascular NRP1 expression has the potential to modify VEGFR2 signalling and alter VEGFA-induced vascular hyperpermeability. Further elucidation of NRP1 expression patterns in different tissues and how the VEGFA/NRP1/VEGFR2 complex presented in *cis* or *trans* can differentially modify cellular signalling will allow for better understanding of how therapeutic targeting of NRP1 may have very different outcomes on vascular function in different tissues engaged in pathological processes.

## Acknowledgements

We thank the BioVis imaging facility, Uppsala University, for expert services. We appreciate expert advice from Dr. Konstantin Gängel, Uppsala University. This study was supported by the Swedish Research Council (2022-00896), the Knut and Alice Wallenberg foundation (KAW 2020.0057 and KAW 2019.0276), Fondation Leducq Transatlantic Network of Excellence Grant in Neurovascular Disease (17 CVD 03) to LCW. MR was supported by Olle Engkvist (218-0057) SSMF (201912) and an EMBO long-term fellowship (ALTF 923–2016).

## Author contributions

Conceptualization, M.R. and L.C.-W.; methodology, M.R. and S.P.; investigation, M.R., S.P., Y.S., E.N.; writing the draft, M.R., S.P. and L.C.-W., final version; all authors; funding acquisition; M.R. and L.C.-W.; resources, L.C.-W.

## Figure legends

**Supplementary Figure 1. NRP1 expression in the ear dermis and back skin of adult mice**

**(A)** Western blot and quantification of NRP1 expression using lung lysates from tamoxifen-treated control *Nrp1^fl/fl^* and *Nrp1* iECKO mice (n = 4).

**(B,C)** Whole mount immunostainings showing NRP1 expression in ear dermis **(B)** and back skin **(C)** of control *Nrp1^fl/fl^*, *Nrp1* iECKO and *Nrp1* iKO mice. Scale bar: 50 μm.

**(D,E)** Images, left, and quantification, right, of vascular and perivascular NRP1 expression in ear skin and back skin of control *Nrp1^fl/lf^* and *Nrp1* iECKO mice. Scale bar: 20 μm.

Error bars; mean ± SEM. Statistical significance used: Two-tailed unpaired Student’s t-test.

**Supplementary Figure 2. Organotypic assessment of VEGFA-induced leakage**

**(A)** Basal permeability of 10 kDa dextran in kidney, back skin, ear dermis, skeletal muscle and heart in control *Nrp1^fl/fl^* and *Nrp1* iECKO mice. Extravasated dextran was measured following extraction from perfused tissues. (n ≥ 8 mice).

**(B,C)** Leakage of 2000 kDa **(B)** and 70 kDa **(C)** dextran after systemic administration of only dextran (PBS) or dextran plus VEGFA in back skin, trachea, kidney, skeletal muscle and heart of wild-type C57Bl/6 mice (n ≥ 3 mice).

**(D,E)** VEGFA-mediated leakage of 70 kDa **(D)** and 2000 kDa **(E)** dextran in kidney, skeletal muscle and heart in control *Nrp1^fl/fl^* and *Nrp1* iECKO mice. Extravasated dextran was measured following extraction from perfused tissues. (n ≥ 8 mice).

Error bars; mean ± SEM. Statistical significance used: Two-tailed unpaired Student’s t-test.

**Supplementary Figure 3. Perivascular NRP1 modifies PLCγ phosphorylation**

**(A-B)** Phosphorylation of PLCγ in response to intradermal PBS or VEGFA injections in the ear dermis of control *Nrp1^fl/fl^* and *Nrp1* iECKO **(A)** or *Nrp1* iKO **(B)** mice. Left, representative images. Right, quantification of phosphorylated PLCγ area per total VEC area normalized to PBS control in the ear dermis of control *Nrp1^fl/fl^* and *Nrp1* iECKO or *Nrp1* iKO mice. n ≥ 3 mice, two or more fields of view / mouse.

**(C-D)** Phosphorylation of PLCγ in response to intradermal PBS or VEGFA injections in the back skin of control *Nrp1^fl/fl^* and *Nrp1* iECKO **(C)** or *Nrp1* iKO **(D)** mice. Left, representative images. Right, quantification of phosphorylated PLCγ area per total VEC area normalized to PBS control in the back skin of control *Nrp1^fl/fl^* and *Nrp1* iECKO or *Nrp1* iKO mice. n ≥ 3 mice, two or more fields of view/ mouse.

Error bars; mean ± SEM. Statistical significance: Two-way ANOVA. Scale bar: 50 μm.

**Video 1: VEGFA-induced leakage in control *Nrp1*^fl/fl^, *Cdh5* Cre-negative littermate mice**

Extravasation of circulating 2000 kDa FITC Dextran (pseudocolor) in control *Nrp1*^fl/fl^, *Cdh5* Cre-negative littermate mice after intradermal injection of VEGFA in the ear dermis. Video corresponds to stills in Figure 1D.

**Video 2: VEGFA-induced leakage in *Nrp1*^fl/fl^; *Cdh5^CreERT2^* (*Nrp1* iECKO) mice**

Extravasation of circulating 2000 kDa FITC Dextran (pseudocolor) in *Nrp1*^fl/fl^; *Cdh5^CreERT2^* (*Nrp1* iECKO) mice after intradermal injection of VEGFA in the ear dermis. Video corresponds to stills in Figure 1D.

**Video 3: VEGFA-induced leakage in control *Nrp1*^fl/fl^, *Actb* Cre-negative littermate mice**

Extravasation of circulating 2000 kDa FITC Dextran (pseudocolor) in control *Nrp1*^fl/fl^, *Actb* Cre-negative littermate mice after intradermal injection of VEGFA in the ear dermis. Video corresponds to stills in Figure 2C.

**Video 4: VEGFA-induced leakage in *Nrp1*^fl/fl^; *Actb^CreERT2^* (*Nrp1* iKO) mice**

Extravasation of circulating 2000 kDa FITC Dextran (pseudocolor) in *Nrp1*^fl/fl^; *Actb^CreERT2^* (*Nrp1* iKO) mice after intradermal injection of VEGFA in the ear dermis. Video corresponds to stills in Figure 2C.

**Video 5: VEGFA-induced leakage in *Nrp1*^fl/fl^; *Cdh5^CreERT2^* (*Nrp1* iECKO) mice treated with isotype control IgG antibody in the ear dermis**

Extravasation of circulating 2000 kDa FITC Dextran (pseudocolor) in *Nrp1*^fl/fl^; *Cdh5^CreERT2^* (*Nrp1* iECKO) mice pre-treated with an isotype control igG antibody and subsequently challenged with intradermal injection of VEGFA in the ear dermis. Video corresponds to stills in Figure 2G.

**Video 6: VEGFA-induced leakage in VEGFR2 ^Y949F/Y949F^; *Nrp1*^fl/fl^; *Cdh5^CreERT2^* (*Nrp1* iECKO) mice treated with a NRP1-VEGFA blocking antibody in the ear dermis**

Extravasation of circulating 2000 kDa FITC Dextran (pseudocolor) in *Nrp1*^fl/fl^; *Cdh5^CreERT2^* (*Nrp1* iECKO) mice pre-treated with a NRP1-VEGFA blocking antibody and subsequently challenged with intradermal injection of VEGFA in the ear dermis. Video corresponds to stills in Figure 2G.

**Video 7: VEGFA-induced leakage in control VEGFR2 ^Y949F/Y949F^; *Nrp1*^fl/fl^; *Cdh5* Cre-negative littermate mice**

Extravasation of circulating 2000 kDa FITC Dextran (pseudocolor) in VEGFR2 ^Y949F/Y949F^; *Nrp1*^fl/fl^; *Cdh5* Cre-negative littermate mice after intradermal injection of VEGFA in the ear dermis. Video corresponds to stills in Figure 4C.

**Video 8: VEGFA-induced leakage in control VEGFR2 ^Y949F/Y949F^; *Nrp1*^fl/fl^; *Cdh5 ^CreERT2^* mice**

Extravasation of circulating 2000 kDa FITC Dextran (pseudocolor) in VEGFR2 ^Y949F/Y949F^; *Nrp1*^fl/fl^; *Cdh5 ^CreERT2^* mice after intradermal injection of VEGFA in the ear dermis. Video corresponds to stills in Figure 4C.

## Methods

### Animals

*In vivo* animal experiments were carried out in accordance with the ethical permit provided by the Committee on the Ethics of Animal Experiments of the University of Uppsala (permit 6789/18). *Nrp1*^fl/fl^ mice contain *loxP* sites flanking exon 2 and were obtained from The Jackson Laboratory (005247). This strain was crossed with *Cdh5*(PAC)-CreER^T2^ mice (kind gift from Dr. Ralf Adams, Max-Planck Institute Münster) or Tmem163Tg(*Actb*-cre)2Mrt mice to generate endothelial-specific *Nrp1* iECKO mice and global *Nrp1* iKO mice, respectively. Mice possessing a knock-in mutation in the VEGFR2 gene, *Kdr*, at the Y949 phosphosite have been described previously (Li et al., 2016). These mice were crossed with *Nrp1* iECKO mice to generate *Vegfr2 ^Y949F/Y949F^; Nrp1* iECKO mice. *Pdgfrb-*GFP reporter mice (Tg(*Pdgfrb*-*EGFP*)jn169Gsat/Mmucd, Stock number 031796-UCD) have been described previously (Jung et al., 2018). Mice were maintained in ventilated cages with group housing (2–5 per cage) and access to water and feed ad libitum. Male and female mice age 8-16 weeks were used. Each experiment was conducted on tissue from at least three age-matched animals representing individual biological repeats (N). To ensure technical reproducibility, where possible multiple images or acquisitions are carried out on each mouse, with the mean of analysed values consituting an individual N. Number of acquired images / movies were chosen to ensure technical reproducibility within ethical bounds. To induce Cre recombinase-mediated gene recombination tamoxifen (SigmaAldrich, T5648) formulated in peanut oil was injected intraperitoneally (80 mg/kg at 4 ml/kg) for 5 consecutive days. The mice were allowed to rest for 4 days before experiments were conducted. For each of the strains used in experiments, control mice were studied in parallel. The exact genotype of control mice is specified in figures and figure legends.

### Intravital vascular leakage assay

Intravital imaging of the mouse ear after intradermal injection of VEGFA has been described previously (Honkura et al., 2018). Briefly, following systemic administration of 2000 kDa FITC (SigmaAldrich, FD2000S) or TRITC Dextran (ThermoFischer Scientific, D7139) by tail-vein injection, mice were sedated by intraperitoneal injection of Ketamine-Xylazine (120 mg/kg Ketamine, 10 mg/kg Xylazine) and the ear secured to a solid support. Mice were maintained at a body temperature of 37 °C for the entire experiment, maximum 90 min. Time-lapse imaging was performed using single-photon microscopy (Leica SP8). For intradermal EC stimulation, a volume of approximately 0.1 μl murine VEGFA164 (Peprotech, 450-32), concentration 100 ng/μl, was injected using a sub-micrometer capillary needle. 10 kDa TRITC Dextran (ThermoFischer Scientific, D1817) was injected with VEGFA to act as a tracer. Leakage sites were identified in time-lapse imaging as defined sites of concentrated dextran in the extravascular space.

### Miles’ assay

Mice were injected intraperitoneally with pyrilamine maleate salt (4 mg/kg, Sigma, P5514) diluted in 0.9% saline, to inhibit histamine release, 30 min prior to tail vein injection of Evans blue (100 μl, 1% Evans blue). Evans blue was allowed to circulate for 10 min before anaesthetising the mice with 3% isoflourane until righting relfexes were lost, switching to 1.5% isoflourane and conducting intradermal injection in the back skin or ear skin of recombinant mouse VEGFA164 (Peprotech, 450-32) (80 ng in 25 μl for back skin, 20ng in 5 μl for the ear skin) or PBS. The skin was excised 30 minutes subsequent to VEGFA injection, placed in formamide over night at 55°C and absorbance at 620 nm was measured using a spectrophotometer and normalized to tissue weight.

### Systemic VEGFA-induced permeability

To assess EC permeability, all tissues were cleaned of excess cartilage, fat and connective tissue. Skeletal muscle was assessed using the tibialis anterior. To analyse VEGFA-induced vascular permeability in the kidney, skeletal muscle and heart, dextran (30 mg/kg) with or without VEGFA (160 μg/kg) were injected systemically through the tail vein. Thirty minutes later mice were anesthetized with Ketamine/Xylazine before intracardiac perfusion with Dulbecco’s phosphate buffered saline (DPBS). Tissues were dissected, washed in DPBS and incubated in formamide for 48 hr at 55 °C. Dextran fluorescence was then measured using a spectrophotometer and normalized to tissue weight. To assess VEGFA-induced leakage microscopically, mixtures of fixable dextran (FITC, Tdb labs; TRITC, ThermoFischer Scientific, D7139) (30 mg/kg) with VEGFA (160 μg/kg) were injected systemically through the tail vein. Thirty minutes later, mice were anesthetized with Ketamine/Xylazine before intracardiac perfusion with DPBS followed by 1% paraformaldehyde (PFA). Tissues were then immersed in 1% PFA for 2 hr before proceeding with immunohistochemistry. For leakage quantification at least three large tile scan areas (≥1 mm^2^) were captured for each mouse.

### Intradermal VEGFA injection in the ear and back skin for immunofluorescent staining

Mice were anesthetised with 3% isoflourane until righting relfexes were lost, switching to 1.5% isoflourane and conducting intradermal injection in the back skin or ear skin of recombinant mouse VEGFA164 (Peprotech, 450-32) or PBS (20 ng VEGFA in 5 μl for the ear dermis and 80 ng VEGFA in 20 μl for the back skin). Five minutes later, mice were anesthetized with Ketamine/Xylazine before intracardiac perfusion with PBS followed by 1% paraformaldehyde (PFA). Tissues were then immersed in 1% PFA for 2 hr before proceeding with immunohistochemistry.

### NRP1 blockade

Anti-NRP1 (R&D systems, AF566) or isotype control IgG (Jackson immunoresearch, 005-000-003MAB006) were administered intradermally in the ear skin (1 µg) or back skin (5 µg), or systemically (2 mg/kg). 24 hours later VEGFA-mediated vascular leakage was assessed using the intravital vascular leakage assay, or through systemic VEGFA delivery in the circulation.

### Immunohistochemistry

Tissues were fixed through intracardiac perfusion with 1% PFA followed by immersion in 1% PFA for 2 hr at room temperature. For whole mount preparation of the ear skin, back skin and trachea, removal of excess cartilage, fat and connective tissue tissues was carried out. Tissues were blocked overnight at 4 °C in Tris-buffered saline (TBS), 5% (w/v) Bovine Serum Albumin (BSA), 0.2% Triton X-100. Samples were incubated overnight with primary antibody in blocking solution, followed by washing in TBS, 0.2% Triton X-100 and incubation with appropriate secondary antibody for 2 hr at room temperature in blocking buffer before washing and mounting in fluorescent mounting medium (DAKO). Images were acquired using a Leica SP8 confocal microscope.

Commercial antibodies used were: rat anti-CD31 (BD Biosciences, 553370), goat anti-CD31 (R&D Systems, AF3628), goat anti-NRP1 (R&D Systems, AF566), chicken anti-GFP (Abcam, Ab13970), rabbit anti-phospho-PLCγ1 (Tyr783) (Cell signaling technology, 2821s), rabbit anti-PLCγ1 (Cell signaling technology, 2822s), and goat anti–VE-cadherin (R&D systems, AF1002). Secondary antibodies against rat (ThermoFischer Scientific; Alexa 488, A21208 and Alexa 594, A21209), rabbit (ThermoFischer Scientific; Alexa 488, A21206 and Alexa 568, A10042), goat (ImmunoResearch Laboratories, Alexa 647, 705-605-147), chicken (ImmunoResearch Laboratories, Alexa 488, 703-545-155) were used. Custom-made pVEC Y685 antibody was prepared by immunizing rabbits with phospho-peptides of the corresponding region in mouse VE-cadherin (New England Peptide) and further verified using immunostaining in Cdh5*^Y685F/Y685F^*mice (Jin et al., 2022). Primary and secondary antibodies were prepared at a dilution of 1:100 and 1:400, respectively unless otherwise stated.

### RNAscope^TM^ *in-situ* hybridization

Detection of NRP1 mRNA in the ear dermis was performed using RNAscope Fluorescent Multiplex Assay (ACD Bio, 322340 and 320851) according to the manufacturer’s instructions (https://www.acdbio.com). Briefly, 10 µm transverse cryosections of the ear dermis and back skin were fixed in chilled 4% PFA for 15 minutes (4°C) then dehydrated by incubating in increasing concentrations of ethanol (50%, 75%, 100%, 100% for 5 minutes each). Following protease IV digest (30 min at RT), NRP1 probe (ACD Bio, 471621) was hybridized on the tissue sections for 2 hr at 40°C. 3-plex negative (ACD Bio, 320871) and positive (ACD Bio, 320811) controls were used to confirm signal specificity. Signal amplification and detection was conducted using TSA-Opal fluorophore (Cy5, 1:1000) (TS-000203). Following the RNAscope procedure, sections were immediately counterstained with Hoechst33342 (Molecular Probes, H3570) (1:1000) and CD31 (AF3628, R&D systems) (1:100) for overnight at 4°C to visualize nuclei and blood vessels, respectively. Images were acquired using a Leica SP8 confocal microscope with at least three sections analysed / mouse. Area of fluorescent dots representing mRNA molecules were quantified and normalized to CD31-positive vessel area.

### Western blot analysis

Lungs from mice were removed and snap frozen. Protein was obtained by mechanical dissociation in RIPA buffer (ThermoFisher Scientific, 89900) supplemented with 50 nM Na_3_VO_4_, Phosphatase inhibitor cocktail (Roche 04906837001) and Protease inhibitor cocktail (Roche, 04693116001). LDS sample buffer (Invitrogen, NP0007) and Sample Reducing Agent (Invitrogen, NP0009) were added to the samples and heated to 70°C for 10 min. Proteins were separated on Nu Page 4–12% Bis-Tris Gel (Invitrogen) in MOPS SDS Running buffer (Invitrogen, NP0001), transferred to PVDF membrane (Thermo scientific, 88518) in NuPAGE transfer buffer (Novex, NP006), 10% methanol and subsequently blocked with 5% BSA in Tris-buffered saline with 0.1% Tween 20 (TBST) for 60 min. The immunoblots were analysed using primary antibodies incubated overnight at 4°C and secondary antibodies linked to horseradish peroxidase (HRP) (Cytiva) incubated for 1 hr at room temperature. After each step filters were washed four times with TBST. HRP signals were visualized by enhanced chemiluminescence (ECL) (Cytiva) (1:25000) and imaged with Chemidoc (Bio-Rad).

Primary antibodies targeting GAPDH (Merck, MAB374), NRP1 (R&D Systems, AF566) were used at a dilution of 1:1000.

### Image quantification

For confocal images, macromolecular leakage was quantified by measurement of tracer area following image thresholding. Tracer area was then normalized to vessel area. For analysis of vascular and peri-vascular NRP1 expression, masks were generated of the CD31 vascular area and the peri-vascular area 3 μm around the vessel (see Figure 3A). NRP1 area following thresholding was quantified within, and normalised to these areas. Phosphorylation values were quantified by measurement of phosphorylated vessel area following image thresholding. Phosphorylation area was then normalized to vessel area. Threshold values were kept constant within experiments.

Analysis of leakage from intravital movies has been described previously (Richards et al., 2021). Briefly, leakage sites were identified in time-lapse imaging as defined sites of concentrated dextran in the extravascular space. To quantify these, their numbers were normalized to vessel length. To assess lag period the time of the appearance of these sites following injection of stimulus was quantified. To assess the extent of barrier disruption the rate of dextran extravasation specifically at these sites was quantified over time. See Honkura et al., 2018 for further details.

All measurements were done with Fiji processing package of Image J2 software.

### Statistical analysis

Data are expressed as mean ± SEM. The statistical tests used were the Students’t test and two-way ANOVA. P-values given are from at least 3 independent samples analysed by two-tailed paired t tests or multiple comparisons obtained from two-way ANOVA. Rate of leakage was compared using linear regression and ANCOVA. All statistical analyses were conducted using GraphPad Prism. A p-value <0.05 was considered statistically significant and significances indicated as p<0.05 (*), p<0.01 (**), and p<0.001 (***). For animal experiments, no statistical methods were used to predetermine sample size. The investigators were blinded to allocation during experiment and outcome assessment.

## References

1. Acevedo, L. M., Barillas, S., Weis, S. M., Göthert, J. R. & Cheresh, D. A. 2008. Semaphorin 3A suppresses VEGF-mediated angiogenesis yet acts as a vascular permeability factor. Blood, 111, 2674–80.

2. Aird, W. C. 2007. Phenotypic heterogeneity of the endothelium: I. Structure, function, and mechanisms. Circ Res, 100, 158–73.

3. Augustin, H. G. & Koh, G. Y. 2017. Organotypic vasculature: From descriptive heterogeneity to functional pathophysiology. Science, 357.

4. Ballmer-Hofer, K., Andersson, A. E., Ratcliffe, L. E. & Berger, P. 2011. Neuropilin-1 promotes VEGFR-2 trafficking through Rab11 vesicles thereby specifying signal output. Blood, 118, 816–26.

5. Bartlett, C. S., Scott, R. P., Carota, I. A., Wnuk, M. L., Kanwar, Y. S., Miner, J. H. & Quaggin, S. E. 2017. Glomerular mesangial cell recruitment and function require the co-receptor neuropilin-1. Am J Physiol Renal Physiol, 313, F1232–F1242.

6. Bayliss, A. L., Sundararaman, A., Granet, C. & Mellor, H. 2020. Raftlin is recruited by neuropilin-1 to the activated VEGFR2 complex to control proangiogenic signaling. Angiogenesis, 23, 371–383.

7. Becker, P. M., Waltenberger, J., Yachechko, R., Mirzapoiazova, T., Sham, J. S., Lee, C. G., Elias, J. A. & Verin, A. D. 2005. Neuropilin-1 regulates vascular endothelial growth factor-mediated endothelial permeability. Circ Res, 96, 1257–65.

8. Bosseboeuf, E., Chikh, A., Chaker, A. B., Mitchell, T. P., Vignaraja, D., Rajendrakumar, R., Khambata, R. S., Nightingale, T. D., Mason, J. C., Randi, A. M., Ahluwalia, A. & Raimondi, C. 2023. Neuropilin-1 interacts with VE-cadherin and TGFBR2 to stabilize adherens junctions and prevent activation of endothelium under flow. Sci Signal, 16, eabo4863.

9. Cai, H. & Reed, R. R. 1999. Cloning and characterization of neuropilin-1-interacting protein: a PSD-95/Dlg/ZO-1 domain-containing protein that interacts with the cytoplasmic domain of neuropilin-1. J Neurosci, 19, 6519–27.

10. Cerani, A., Tetreault, N., Menard, C., Lapalme, E., Patel, C., Sitaras, N., Beaudoin, F., Leboeuf, D., DE Guire, V., Binet, F., Dejda, A., Rezende, F. A., Miloudi, K. & Sapieha, P. 2013. Neuron-derived semaphorin 3A is an early inducer of vascular permeability in diabetic retinopathy via neuropilin-1. Cell Metab, 18, 505–18.

11. Corti, F., Ristori, E., Rivera-Molina, F., Toomre, D., Zhang, J., Mihailovic, J., Zhuang, Z. W. & Simons, M. 2022. Syndecan-2 selectively regulates VEGF-induced vascular permeability. Nature Cardiovascular Research, 1, 518–528.

12. Fantin, A., Lampropoulou, A., Senatore, V., Brash, J. T., Prahst, C., Lange, C. A., Liyanage, S. E., Raimondi, C., Bainbridge, J. W., Augustin, H. G. & Ruhrberg, C. 2017. VEGF165-induced vascular permeability requires NRP1 for ABL-mediated SRC family kinase activation. J Exp Med, 214, 1049–1064.

13. Fernández-Robredo, P., Selvam, S., Powner, M. B., Sim, D. A. & Fruttiger, M. 2017. Neuropilin 1 Involvement in Choroidal and Retinal Neovascularisation. PLoS One, 12, e0169865.

14. Gioelli, N., Neilson, L. J., Wei, N., Villari, G., Chen, W., Kuhle, B., Ehling, M., Maione, F., Willox, S., Brundu, S., Avanzato, D., Koulouras, G., Mazzone, M., Giraudo, E., Yang, X. L., Valdembri, D., Zanivan, S. & Serini, G. 2022. Neuropilin 1 and its inhibitory ligand mini-tryptophanyl-tRNA synthetase inversely regulate VE-cadherin turnover and vascular permeability. Nat Commun, 13, 4188.

15. Goshima, Y., Hori, H., Sasaki, Y., Yang, T., Kagoshima-Maezono, M., Li, C., Takenaka, T., Nakamura, F., Takahashi, T., Strittmatter, S. M., Misu, Y. & Kawakami, T. 1999. Growth cone neuropilin-1 mediates collapsin-1/Sema III facilitation of antero- and retrograde axoplasmic transport. J Neurobiol, 39, 579–89.

16. Gu, C., Limberg, B. J., Whitaker, G. B., Perman, B., Leahy, D. J., Rosenbaum, J. S., Ginty, D. D. & Kolodkin, A. L. 2002. Characterization of neuropilin-1 structural features that confer binding to semaphorin 3A and vascular endothelial growth factor 165. J Biol Chem, 277, 18069–76.

17. Hong, T. M., Chen, Y. L., Wu, Y. Y., Yuan, A., Chao, Y. C., Chung, Y. C., Wu, M. H., Yang, S. C., Pan, S. H., Shih, J. Y., Chan, W. K. & Yang, P. C. 2007. Targeting neuropilin 1 as an antitumor strategy in lung cancer. Clin Cancer Res, 13, 4759–68.

18. Honkura, N., Richards, M., Lavina, B., Sainz-Jaspeado, M., Betsholtz, C. & Claesson-Welsh, L. 2018. Intravital imaging-based analysis tools for vessel identification and assessment of concurrent dynamic vascular events. Nat Commun, 9, 2746.

19. Horowitz, A. & Seerapu, H. R. 2012. Regulation of VEGF signaling by membrane traffic. Cell Signal, 24, 1810–20.

20. Hu, B., Guo, P., Bar-Joseph, I., Imanishi, Y., Jarzynka, M. J., Bogler, O., Mikkelsen, T., Hirose, T., Nishikawa, R. & Cheng, S. Y. 2007. Neuropilin-1 promotes human glioma progression through potentiating the activity of the HGF/SF autocrine pathway. Oncogene, 26, 5577–86.

21. Jin, Y., Ding, Y., Richards, M., Kaakinen, M., Giese, W., Baumann, E., Szymborska, A., Rosa, A., Nordling, S., Schimmel, L., Akmeriç, E. B., Pena, A., Nwadozi, E., Jamalpour, M., Holstein, K., Sáinz-Jaspeado, M., Bernabeu, M. O., Welsh, M., Gordon, E., Franco, C. A., Vestweber, D., Eklund, L., Gerhardt, H. & Claesson-Welsh, L. 2022. Tyrosine-protein kinase Yes controls endothelial junctional plasticity and barrier integrity by regulating VE-cadherin phosphorylation and endocytosis. Nature Cardiovascular Research, 1, 1156–1173.

22. Jung, B., Arnold, T. D., Raschperger, E., Gaengel, K. & Betsholtz, C. 2018. Visualization of vascular mural cells in developing brain using genetically labeled transgenic reporter mice. J Cereb Blood Flow Metab, 38, 456–468.

23. Kawakami, T., Tokunaga, T., Hatanaka, H., Kijima, H., Yamazaki, H., Abe, Y., Osamura, Y., Inoue, H., Ueyama, Y. & Nakamura, M. 2002. Neuropilin 1 and neuropilin 2 co-expression is significantly correlated with increased vascularity and poor prognosis in nonsmall cell lung carcinoma. Cancer, 95, 2196–201.

24. Kawasaki, T., Kitsukawa, T., Bekku, Y., Matsuda, Y., Sanbo, M., Yagi, T. & Fujisawa, H. 1999. A requirement for neuropilin-1 in embryonic vessel formation. Development, 126, 4895–902.

25. Kitsukawa, T., Shimono, A., Kawakami, A., Kondoh, H. & Fujisawa, H. 1995. Overexpression of a membrane protein, neuropilin, in chimeric mice causes anomalies in the cardiovascular system, nervous system and limbs. Development, 121, 4309–18.

26. Koch, S., Van Meeteren, L. A., Morin, E., Testini, C., Weström, S., Björkelund, H., Le Jan, S., Adler, J., Berger, P. & Claesson-Welsh, L. 2014. NRP1 presented in trans to the endothelium arrests VEGFR2 endocytosis, preventing angiogenic signaling and tumor initiation. Dev Cell, 28, 633–46.

27. Lanahan, A., Zhang, X., Fantin, A., Zhuang, Z., Rivera-Molina, F., Speichinger, K., Prahst, C., Zhang, J., Wang, Y., Davis, G., Toomre, D., Ruhrberg, C. & Simons, M. 2013. The neuropilin 1 cytoplasmic domain is required for VEGF-A-dependent arteriogenesis. Dev Cell, 25, 156–68.

28. Lanahan, A. A., Hermans, K., Claes, F., Kerley-Hamilton, J. S., Zhuang, Z. W., Giordano, F. J., Carmeliet, P. & Simons, M. 2010. VEGF receptor 2 endocytic trafficking regulates arterial morphogenesis. Dev Cell, 18, 713–24.

29. Li, X., Padhan, N., Sjöström, E. O., Roche, F. P., Testini, C., Honkura, N., Sáinz-Jaspeado, M., Gordon, E., Bentley, K., Philippides, A., Tolmachev, V., Dejana, E., Stan, R. V., Vestweber, D., Ballmer-Hofer, K., Betsholtz, C., Pietras, K., Jansson, L. & Claesson-Welsh, L. 2016. VEGFR2 pY949 signalling regulates adherens junction integrity and metastatic spread. Nat Commun, 7, 11017.

30. Mamluk, R., Gechtman, Z., Kutcher, M. E., Gasiunas, N., Gallagher, J. & Klagsbrun, M. 2002. Neuropilin-1 binds vascular endothelial growth factor 165, placenta growth factor-2, and heparin via its b1b2 domain. J Biol Chem, 277, 24818–25.

31. Matsumoto, T., Bohman, S., Dixelius, J., Berge, T., Dimberg, A., Magnusson, P., Wang, L., Wikner, C., Qi, J. H., Wernstedt, C., Wu, J., Bruheim, S., Mugishima, H., Mukhopadhyay, D., Spurkland, A. & Claesson-Welsh, L. 2005. VEGF receptor-2 Y951 signaling and a role for the adapter molecule TSAd in tumor angiogenesis. Embo j, 24, 2342–53.

32. Mcgowan, S. E. & Mccoy, D. M. 2021. Neuropilin-1 directs PDGFRalpha-entry into lung fibroblasts and signaling from very early endosomes. Am J Physiol Lung Cell Mol Physiol, 320, L179–L192.

33. Miao, H. Q., Lee, P., Lin, H., Soker, S. & Klagsbrun, M. 2000. Neuropilin-1 expression by tumor cells promotes tumor angiogenesis and progression. FASEB J, 14, 2532–9.

34. Morin, E., Lindskog, C., Johansson, M., Egevad, L., Sandström, P., Harmenberg, U., Claesson-Welsh, L. & Sjöberg, E. 2020. Perivascular Neuropilin-1 expression is an independent marker of improved survival in renal cell carcinoma. The Journal of Pathology, 250, 387–396.

35. Morin, E., Sjöberg, E., Tjomsland, V., Testini, C., Lindskog, C., Franklin, O., Sund, M., Öhlund, D., Kiflemariam, S., Sjöblom, T. & Claesson-Welsh, L. 2018. VEGF receptor-2/neuropilin 1 trans-complex formation between endothelial and tumor cells is an independent predictor of pancreatic cancer survival. The Journal of Pathology, 246, 311–322.

36. Naccache, S. N., Hasson, T. & Horowitz, A. 2006. Binding of internalized receptors to the PDZ domain of GIPC/synectin recruits myosin VI to endocytic vesicles. Proc Natl Acad Sci U S A, 103, 12735–40.

37. Orsenigo, F., Giampietro, C., Ferrari, A., Corada, M., Galaup, A., Sigismund, S., Ristagno, G., Maddaluno, L., Koh, G. Y., Franco, D., Kurtcuoglu, V., Poulikakos, D., Baluk, P., Mcdonald, D., Grazia Lampugnani, M. & Dejana, E. 2012. Phosphorylation of VE-cadherin is modulated by haemodynamic forces and contributes to the regulation of vascular permeability in vivo. Nat Commun, 3, 1208.

38. Pan, Q., Chanthery, Y., Liang, W. C., Stawicki, S., Mak, J., Rathore, N., Tong, R. K., Kowalski, J., Yee, S. F., Pacheco, G., Ross, S., Cheng, Z., LE Couter, J., Plowman, G., Peale, F., Koch, A. W., Wu, Y., Bagri, A., Tessier-Lavigne, M. & Watts, R. J. 2007. Blocking neuropilin-1 function has an additive effect with anti-VEGF to inhibit tumor growth. Cancer Cell, 11, 53–67.

39. Ponuwei, G. A. 2016. A glimpse of the ERM proteins. J Biomed Sci, 23, 35.

40. Raimondi, C., Fantin, A., Lampropoulou, A., Denti, L., Chikh, A. & Ruhrberg, C. 2014. Imatinib inhibits VEGF-independent angiogenesis by targeting neuropilin 1-dependent ABL1 activation in endothelial cells. J Exp Med, 211, 1167–83.

41. Richards, M., Nwadozi, E., Pal, S., Martinsson, P., Kaakinen, M., Gloger, M., Sjöberg, E., Koltowska, K., Betsholtz, C., Eklund, L., Nordling, S. & Claesson-Welsh, L. 2022. Claudin5 protects the peripheral endothelial barrier in an organ and vessel-type-specific manner. Elife, 11.

42. Richards, M., Pal, S., Sjöberg, E., Martinsson, P., Venkatraman, L. & Claesson-Welsh, L. 2021. Intra-vessel heterogeneity establishes enhanced sites of macromolecular leakage downstream of laminin α5. Cell Rep, 35, 109268.

43. Roth, L., Prahst, C., Ruckdeschel, T., Savant, S., Weström, S., Fantin, A., Riedel, M., Héroult, M., Ruhrberg, C. & Augustin, H. G. 2016. Neuropilin-1 mediates vascular permeability independently of vascular endothelial growth factor receptor-2 activation. Sci Signal, 9, ra42.

44. Sarris, M., Andersen, K. G., Randow, F., Mayr, L. & Betz, A. G. 2008. Neuropilin-1 expression on regulatory T cells enhances their interactions with dendritic cells during antigen recognition. Immunity, 28, 402–13.

45. Shahrabi-Farahani, S., Gallottini, M., Martins, F., Li, E., Mudge, D. R., Nakayama, H., Hida, K., Panigrahy, D., D’amore, P. A. & Bielenberg, D. R. 2016. Neuropilin 1 Receptor Is Up-Regulated in Dysplastic Epithelium and Oral Squamous Cell Carcinoma. Am J Pathol, 186, 1055–64.

46. Sherafat, A., Pfeiffer, F., Reiss, A. M., Wood, W. M. & Nishiyama, A. 2021. Microglial neuropilin-1 promotes oligodendrocyte expansion during development and remyelination by trans-activating platelet-derived growth factor receptor. Nat Commun, 12, 2265.

47. Sjoberg, E., Melssen, M., Richards, M., Ding, Y., Chanoca, C., Chen, D., Nwadozi, E., Pal, S., Love, D. T., Ninchoji, T., Shibuya, M., Simons, M., Dimberg, A. & Claesson-Welsh, L. 2023. Endothelial VEGFR2-PLCgamma signaling regulates vascular permeability and antitumor immunity through eNOS/Src. J Clin Invest, 133.

48. Soker, S., Miao, H. Q., Nomi, M., Takashima, S. & Klagsbrun, M. 2002. VEGF165 mediates formation of complexes containing VEGFR-2 and neuropilin-1 that enhance VEGF165-receptor binding. J Cell Biochem, 85, 357–68.

49. Soker, S., Takashima, S., Miao, H. Q., Neufeld, G. & Klagsbrun, M. 1998. Neuropilin-1 is expressed by endothelial and tumor cells as an isoform-specific receptor for vascular endothelial growth factor. Cell, 92, 735–45.

50. Sun, Z., Li, X., Massena, S., Kutschera, S., Padhan, N., Gualandi, L., Sundvold-Gjerstad, V., Gustafsson, K., Choy, W. W., Zang, G., Quach, M., Jansson, L., Phillipson, M., Abid, M. R., Spurkland, A. & Claesson-Welsh, L. 2012. VEGFR2 induces c-Src signaling and vascular permeability in vivo via the adaptor protein TSAd. J Exp Med, 209, 1363–77.

51. Termini, C. M., Pang, A., Fang, T., Roos, M., Chang, V. Y., Zhang, Y., Setiawan, N. J., Signaevskaia, L., Li, M., Kim, M. M., Tabibi, O., Lin, P. K., Sasine, J. P., Chatterjee, A., Murali, R., Himburg, H. A. & Chute, J. P. 2021. Neuropilin 1 regulates bone marrow vascular regeneration and hematopoietic reconstitution. Nat Commun, 12, 6990.

52. Wang, J., Wang, S., Li, M., Wu, D., Liu, F., Yang, R., Ji, S., Ji, A. & Li, Y. 2015. The Neuropilin-1 Inhibitor, ATWLPPR Peptide, Prevents Experimental Diabetes-Induced Retinal Injury by Preserving Vascular Integrity and Decreasing Oxidative Stress. PLoS One, 10, e0142571.

53. Wessel, F., Winderlich, M., Holm, M., Frye, M., Rivera-Galdos, R., Vockel, M., Linnepe, R., Ipe, U., Stadtmann, A., Zarbock, A., Nottebaum, A. F. & Vestweber, D. 2014. Leukocyte extravasation and vascular permeability are each controlled in vivo by different tyrosine residues of VE-cadherin. Nat Immunol, 15, 223–30.

54. Wnuk, M., Anderegg, M. A., Graber, W. A., Buergy, R., Fuster, D. G. & Djonov, V. 2017. Neuropilin1 regulates glomerular function and basement membrane composition through pericytes in the mouse kidney. Kidney Int, 91, 868–879.

