## Supplementary figures and images for "Neuropilin-1 controls vascular permeability through juxtacrine regulation of endothelial adherens junctions"

### Supplemental figures

#### Supplemental figure 1

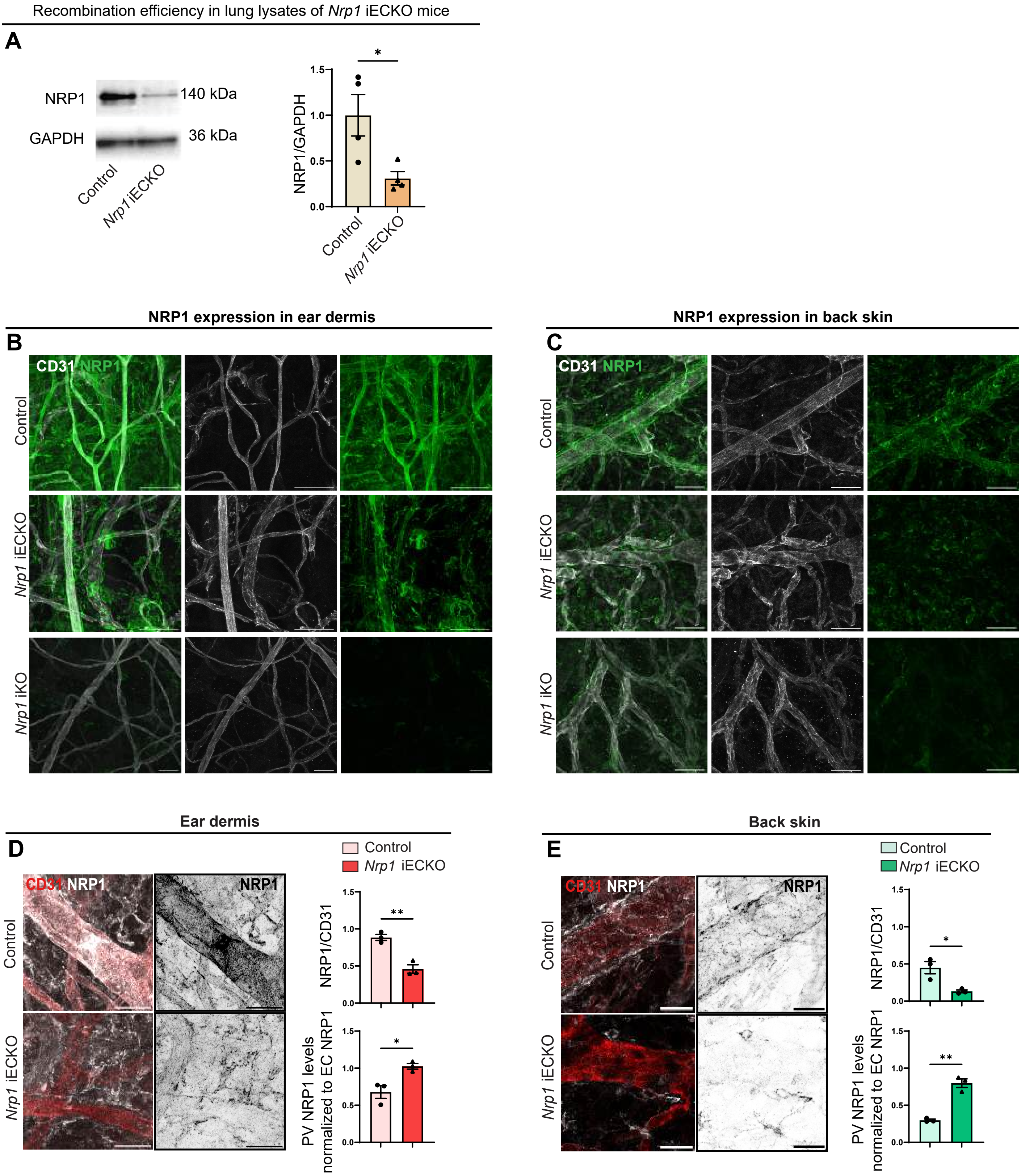

#### Supplemental figure 2

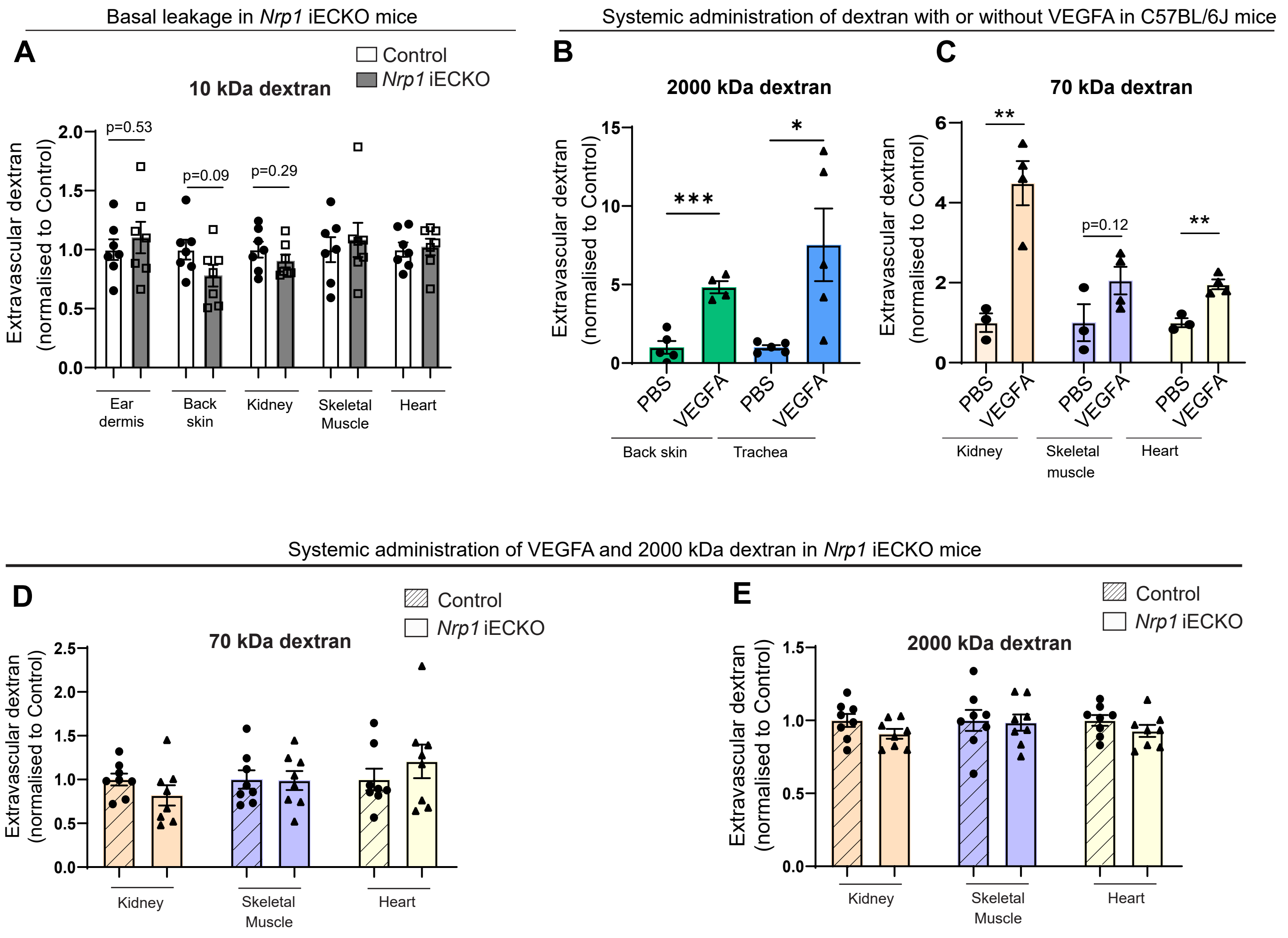

#### Supplemental figure 3

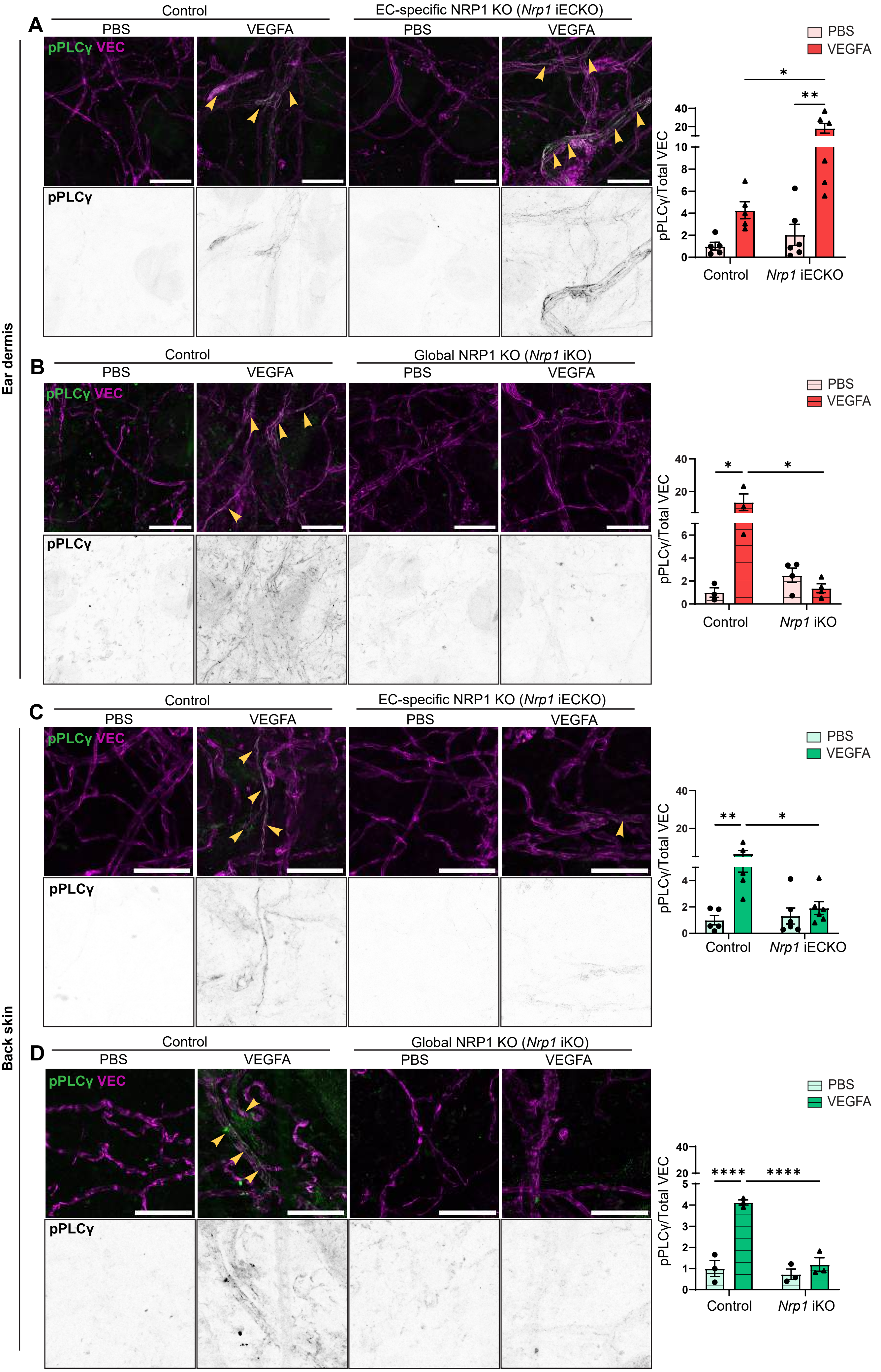
